# A novel hybrid protein promotes Aβ clearance and reduces inflammatory response through MerTK

**DOI:** 10.1101/2021.11.03.467048

**Authors:** Lorena P. Samentar, Arnold Salazar, Pei-Pei Pan, Kayvon Etebar, Kelly Choy, Durin Uddin, Pauline Eliseeff, Adrienne Marrie Bugayong, Jose Antonio Ma. G. Garrido, Aurora Emini, Nicole Rock, Nora Blanca Caberoy

## Abstract

Alzheimer’s disease (AD) is the world’s leading cause of dementia and the most common neurodegenerative disorder. Its major pathological features are amyloid beta (Aβ) plaques, tau tangles, and neuroinflammation that eventually leads to massive death of nerve cells. Even with the multifactorial aspect of AD, the most accepted theory is that Aβ is the driving force of AD pathogenesis. We engineered a novel hybrid protein that facilitates the phagocytosis of Aβ and redirect its clearance to the noninflammatory Mer tyrosine kinase (MerTK) pathway. The novel hybrid protein facilitates robust uptake and clearance of Aβ in BV2 microglia through MerTK receptor with reduced production of inflammatory factors and oxidative products. In APP/PS1 transgenic AD mouse model, intraperitoneal administration of the hybrid protein for two months results in significant reduction of Aβ burden in the brain and protection of nerve cells from dying. Taken together, our results suggest that the novel hybrid may have the potential for AD treatment by targeting both Aβ clearance and reduction of inflammation.

## INTRODUCTION

Alzheimer’s disease (AD) is a chronic progressive neurodegenerative disorder characterized by cognitive dysfunction, psychiatric symptoms, behavioral disturbances, and difficulties with performing activities of daily living (Burns and Iliffe, 2009) that ultimately leads to dementia and death (Bertram and Tanzi, 2019). The disease affects about 5.8M Americans (Alzheimer’s Association, 2020) and nearly 48M people worldwide (Du et al, 2018). AD dementia is projected to have a devastating impact on global populations by 2050 when it is predicted to affect about 131 million people (Cummings et al., 2020). Because AD is a multifactorial disease, the exact pathophysiology is poorly understood. However, the most accepted theory is that the presence of aggregated Aβ initiates downstream effects, including tau hyperphosphorylation, chronic neuroinflammation, oxidative stress, and loss of synapses and neurons that ultimately lead to considerable brain atrophy and cognitive decline (Bertram and Tanzi, 2019; Du et al., 2018); Spangenberg and Green, 2017; Butterfield and Lauderback, 2002). The Aβ hypothesis proposes that age-dependent accumulation of the Aβ peptide is a critical step in the cascade of pathological events that cause deficits in synaptic functionality and lead to severe neurodegeneration in AD (Bohrmann et al., 2012). The exact mechanism of how Aβ in AD leads to synaptic loss and neuronal death remains to be elucidated.

Microglia are the brain’s resident immune cells. They function as the brain’s first line of defense to protect the central nervous system from injury and invading pathogens. One of the more extensively studied functions of microglia in the brain is their role in clearance via phagocytosis. As a result, these cells both protect the brain from invading pathogens as well as remove cellular debris from the neural environment (Spangenberg and Green, 2017). In AD, microglial cells play an important role in disease progression by clearing Aβ deposits and initiating phagocytic activity (Wolfe, 2016; Yan et al., 2012; Yu and Ye, 2015; Stalder et al., 1999; Pan et al., 2011). Aβ phagocytosis is primarily mediated by pattern recognition receptors (PRRs) including Receptor for Advanced Glycation End-products (RAGE), among others (Yan et al., 2012; Gąsiorowski et al., 2018; Fang et al.,2010; Fang et al., 2018; Derk et al., 2018). Microglial phagocytosis of Aβ through RAGE leads to increased production of proinflammatory mediators (Behl, 2017) and microglia migration/infiltration, which increases neuroinflammation and neuronal damage (Yan et al., 2012; Dong et al., 2019). Inhibition of RAGE suppresses microglia activation and the associated inflammatory response while overexpression of RAGE in microglia increases tissue infiltration of glial cells, microglia activation, Aβ accumulation, and deterioration of cognitive functions (Yu and Ye, 2015; Fang et al., 2018). Microglia-mediated neuroinflammation is one of the most remarkable hallmarks in neurodegeneration (Wolfe, 2016) because the inflammatory process can directly damage neurons while concurrently promoting protein aggregations (Dong et al., 2019). Microglial cells activated by Aβ not only induce the expression of proinflammatory cytokines including interleukin (IL)-1β, IL-6, IL-8, and tumor necrosis factor-α (TNF-α), but also of chemokines and reactive oxygen and nitrogen species, all of which cause neuronal damage (Bertram and Tanzi, 2019; Wolfe, 2016; Caberoy et al, 2012a)

MerTK, a member of the Tyro-Axl-MerTK (TAM) family of receptor tyrosine kinases, is a macrophage receptor that mediates the binding and phagocytosis of apoptotic cells, a process known as efferocytosis (Lemke, 2013; Lemke and Rothlin, 2008). MerTK interacts with apoptotic cells through the bridging molecules Gas6 or protein S (Cai et al., 2018), Tulp1, Tubby and Gal3 (Caberoy et al., 2010a; Caberoy et al., 2012a; Caberoy et al., 2012b). MerTK signaling in macrophages promotes ERK activation and the resolution of inflammation by promoting the synthesis of inflammation resolution mediators (Lemke, 2013; Lemke and C. V. Rothlin, 2008; Rothlin et al., 2007). Loss of these responses by genetic targeting of MerTK in mice can lead to chronic diseases of inflammation and impaired resolution (Cai et al., 2018). MerTK also plays an important role in inhibition of Toll-like receptor (TLR)-mediated innate immune response by activating STAT1, which selectively induces production of suppressors of cytokine signaling SOCS1 and SOCS3 (Lemke, 2013; Lee and Chun, 2019; Zizzo and Cohen, 2018). In mouse microglial cells, MerTK is involved in the inhibition of Toll-like receptor-mediated inflammatory pathways in the presence of its ligand Gas6 (Gilchrist et al., 2020).

AD is progressively debilitating, inevitability lethal, and astronomically expensive but until now, it has no effective treatment (Wolfe, 2016). Most available treatments only address the cognitive and behavioral symptoms (Dong et al., 2019; Cummings et al., 2018) but do not really delay or stop the death of nerve cells that cause the symptoms and make Alzheimer’s fatal. The only exception is aducanumab (Aduhelm) as the first novel therapy approved for AD since 2003 and the only therapy directed at the underlying pathophysiology of AD (Xu et al., 2018). Hence, therapies preventing, delaying the onset, slowing the progression, and improving the symptoms of AD are still urgently needed (Xu et al., 2018). The use of hybrid molecules that feature two or more bioactive components has recently gained momentum in drug design and development to simultaneously modulate multiple drug targets of multifactorial diseases (Bérub, 2016; Decker, 2012; Ibrar et al., 2018; Kaur and Silakari, 2018) like AD. The AD drug discovery pipeline has expanded from Aβ and tau targets to also include inflammation, synapse and neuronal protection, cellular stress, lipid metabolism, and epigenetic intervention (Xu et al., 2018). Being a multifactorial disease, AD treatment most likely will require combination therapies. Hence, this study with the novel hybrid protein that we engineered targets both Aβ clearance and reduction of inflammation.

The proposed mechanism of action of our novel therapeutic strategy has an inherent advantage because Aβ clearance is directed to a noninflammatory receptor, thereby reducing not only the Aβ burden but also the massive inflammation that leads to the death of nerve cells in AD. While the use of hybrid molecules that feature two or more potentially bioactive components has been used in drug development to simultaneously modulate multiple drug targets of multifactorial diseases (Ibrar et al., 2018), to our knowledge, this is the first research that makes use of hybrid molecule to target the phagocytic clearance of a pathogenic molecule through a specific and noninflammatory receptor. Moreover, despite our knowledge of the varied roles of MerTK receptor affecting health and diseases, modulation or use of this receptor has not been targeted for phagocytosis-based therapy using customized ligands. Our approach can highly impact translational research as this can easily be adapted to the development of therapeutics to divert the clearance of harmful molecules through precise targeting of the ligand to a specific cellular receptor to minimize off target effects.

## RESULTS

### Generation and characterization of the novel hybrid protein

Previously, we have identified Tubby protein with its minimal phagocytic domain (MPD) that facilitates the phagocytosis of cellular debris in the retina through MerTK (Caberoy et al., 2010a). Unlike RAGE and other PRRs, phagocytosis through MerTK is considered silent because it does not result in an inflammatory response (Lemke, 2013; Lemke and Rothlin, 2008; Caberoy et al., 2010a; Caberoy et al. 2012a; Rothlin et al., 2007). In view of developing an Alzheimer’s therapy, our objective is to divert the clearance of Aβ from inflammatory RAGE to the non-inflammatory MerTK pathway. Hence, we created a hybrid molecule that can bind to both MerTK and Aβ. This hybrid protein contained the minimal phagocytic domain (MPD) of Tubby that can recognize MerTK and the Aβ binding peptide (AβBP) that can specifically bind to Aβ (Figure 1A). The idea is that the hybrid protein will act as molecular bridge with the AβBP binding to Aβ and the MPD of Tubby recognizing MerTK so that together they will redirect Aβ clearance from RAGE to MerTK.

**Figure 1.**
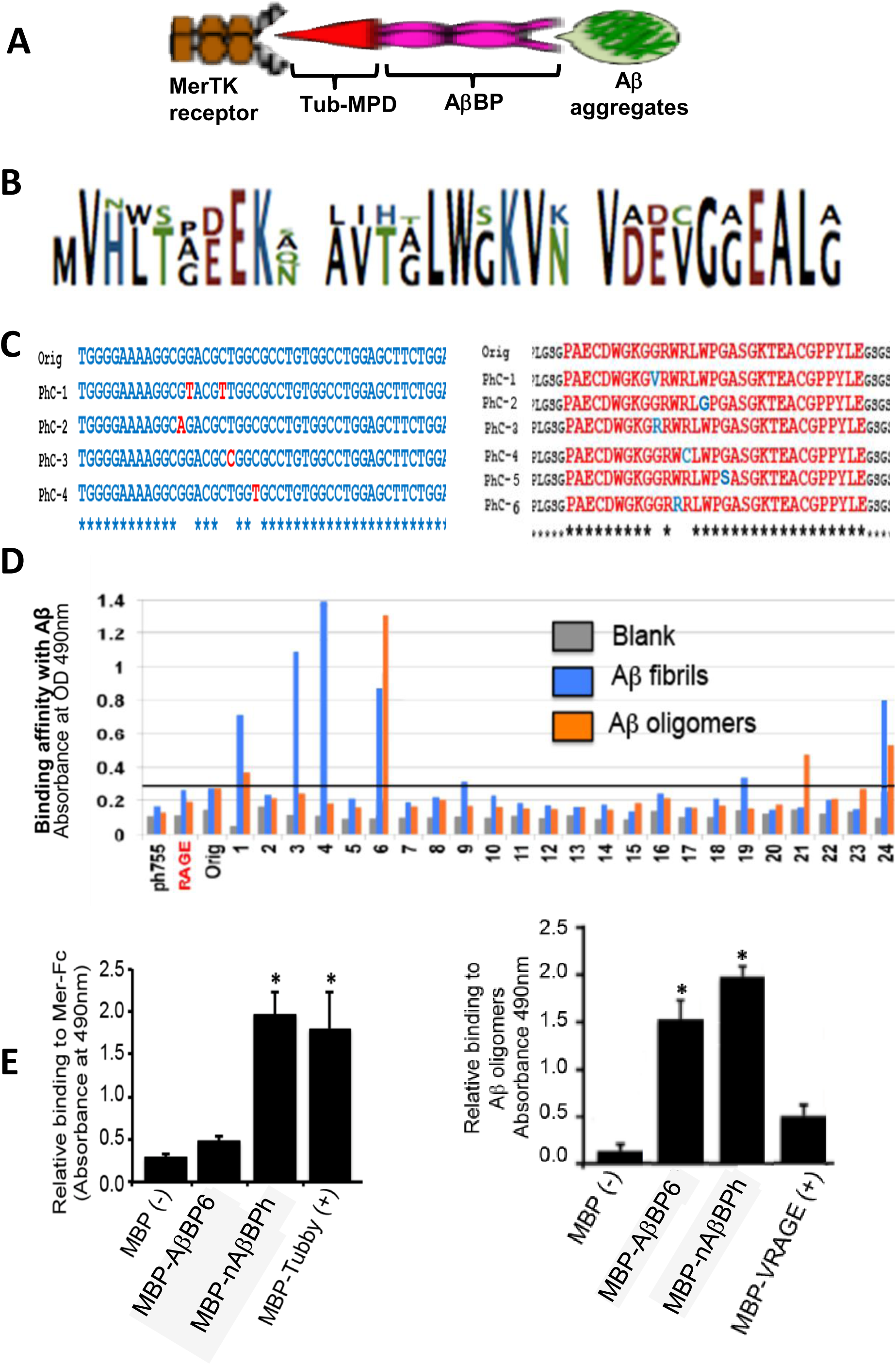
Generation and characterization of novel AβBP hybrid (nAβBPh) with high binding affinity to Aβ. (A) Schematic diagram of the hybrid protein. (B) The 24-aa AβBP-containing consensus sequence was generated by aligning the 128 known Aβ-binding proteins in the literature using the NCBI-COBALT software (Papadopoulos and Agarwala, 2007). (C) Analyses of the nucleotide (left) and amino acid sequences (right) of mutant AβBP phage clones showed transition and transversion mutations resulting to changes in the amino acid sequences. (D) The binding affinities of the evolved AβBPs to Aβ oligomers and fibrils were determined using ELISA. AβBP6 that showed high binding to Aβ oligomers and fibrils was chosen and fused to the MPD of Tubby to create nAβBPh. The empty vector, and vectors with coding regions of RAGE and the AβBP consensus sequence were included as controls. (E) The ability of the hybrid protein to bind to both MerTK (left) and Aβ oligomers (right) was analyzed using ELISA. Note that nAβBPh binds to Mer-Fc, the soluble ligand binding domain of MerTK receptor, with the same efficiency as full-length Tubby even if only the MPD part of Tubby was used. Also, the nAβBPh binds to Aβ oligomers similar to AβBP6, and 4X better than the variable domain of RAGE (VRAGE) where the ligand binds. Data are expressed as mean ± standard error of the mean (SEM). P < 0.05, Student’s t test; n = 3 independent experiments.

Since the MPD sequence that is recognized by MerTK is already known, we generated the Aβ binding peptide (AβBP) consensus sequence using a bioinformatics approach for the creation of the hybrid protein (Figure 1A). We aligned about 128 known sequences of Aβ-binding proteins in the literature using the NCBI-COBALT (Papadopoulos and Agarwala, 2007) software to come up with a 24-aa AβBP consensus sequence (Figure 1B). Next, we subjected the AβBP consensus sequence to *in vitro* protein evolution by PCR mutagenesis and open reading frame dual phage display (Caberoy et al., 2010b; Caberoy et al., 2009). Mutant AβBP phage clones show transition and transversion mutations (Figure 1C left) that result to changes in the amino acid sequences (Figure 1C right). The evolved AβBPs had varying binding affinities to Aβ oligomers and fibrils (Figure 1D), but AβBP6 showed high binding to Aβ fibrils and the highest binding to Aβ oligomers. Therefore, we chose AβBP6 and generated the novel hybrid protein (**nAβBPh**) by fusing two repeats each of AβBP6 and the MPD of Tubby (Figure 1A).

For the hybrid protein to perform as designed, it should bind to both MerTK and Aβ and it should be stable under physiological conditions. We demonstrated through ELISA that the hybrid protein bound to Mer-Fc (the extracellular domain of MerTK), with the same efficiency as full-length Tubby even if only the MPD part of Tubby was used (Figure 1E left), and to Aβ oligomers similar to AβBP6 and 4X better than the variable domain of RAGE (VRAGE) (Figure 1E right). Protein aggregation assays showed that at pH 7.0 and at 8mg/mL concentration (the average working concentration that we used), the hybrid protein was stable. It aggregated only at 70°C (Supplementary Figure 1).

### The hybrid protein facilitates significantly higher phagocytic efficiency of Aβ by BV2 cells

The BV2 mouse microglial cell line derived from C57/Bl6 murine microglia immortalized by v-raf/v-myc carrying J2 retrovirus (Blasi et al., 1990) is one of the most popular cellular models to study neuroinflammation and neurodegeneration because of its phagocytic capabilities and responsiveness not only to lipopolysaccharide (LPS) but also to Aβ (Stansley, et al., 2012; Timmerman et al., 2018). Given that the hybrid protein could bind to both MerTK and Aβ, we tested whether it could facilitate phagocytosis of Aβ by BV2 cells. At a concentration dependent manner, the hybrid protein facilitated significantly higher phagocytosis efficiency of BV2 cells in terms of both the internalization of Aβ and the percentage of phagocytic cells (Figure 2A to Figure 2C). Complementary to confocal microscopy, we performed fluorescence activated cell sorting (FACS) as an independent quantification of phagocytosed Aβ. Consistent with the confocal results, the hybrid protein facilitated robust phagocytosis of both Aβ oligomers and fibrils (Supplementary Figure 3).

**Figure 2.**
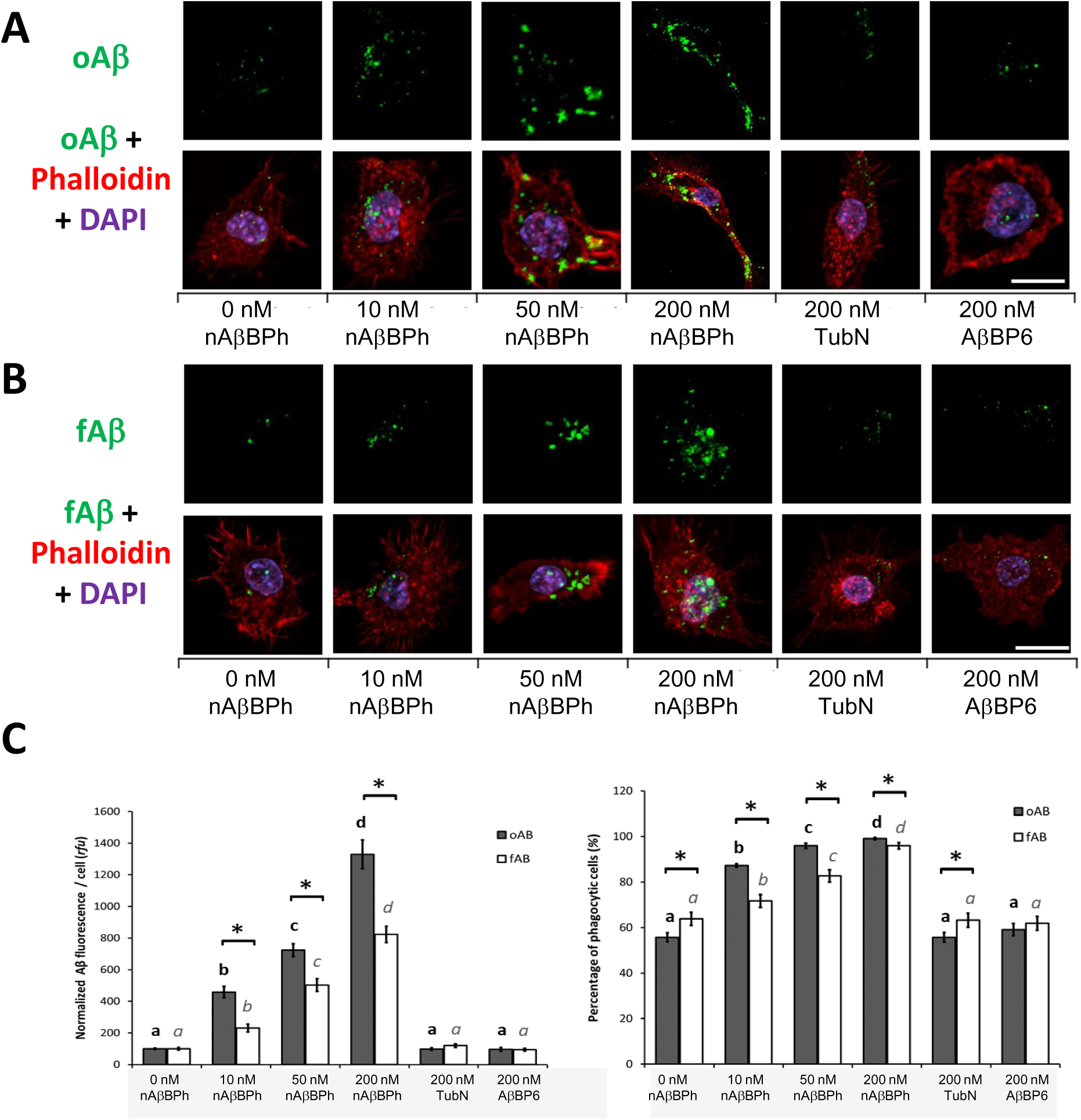
The hybrid protein facilitates significantly higher phagocytic efficiency of Aβ by BV2 cells. (A) Representative confocal images for oAβ, and (B) fAβ. (C) Quantification using ImageJ Fiji showed that phagocytosis efficiency of BV2 cells in terms of the fluorescence signal of internalized Aβ (left) and the percentage of phagocytic cells (right) significantly increased in the presence of the hybrid protein at a concentration dependent manner. The fluorescence intensity and exposure settings were kept constant. Fluorescence data were normalized to the negative control (0 nM nAβBPh) following background subtraction using cells not fed with Aβ. Scale bar = 10 μm. n = 10 fields with at least 20 cells/field (source data **Supplementary Figure 2**). Correlation coefficients between Aβ fluorescence and percentage of phagocytic cells in Aβ oligomers = 0.92 and in Aβ fibrils = 0.94. Different letters denote statistical significance at P ≤ 0.05 between treatments using unpaired, two-tailed Student’s t-test. Asterisk (*) denotes significant differences between the means of oAβ and fAβ in the same treatment. Data are expressed as mean ± SEM.

In view of the utility of the hybrid protein for AD treatment, we next tested whether it can also facilitate the uptake of physiologically generated Aβ through an *ex vivo* phagocytosis assay following Borhmann et al. (2012). We used serial sections of AD mouse brain previously verified to have Aβ plaques (Figure 3A). In the presence of the hybrid protein, GFP-expressing BV2 cells are observed not only to have internalized more Aβ but also to congregate around Aβ plaques in APP/PS1 AD mouse brain (Figure 3B). All these observations show a robust hybrid protein-facilitated phagocytosis of both synthetic and physiologically generated Aβ, and its eventual degradation.

**Figure 3.**
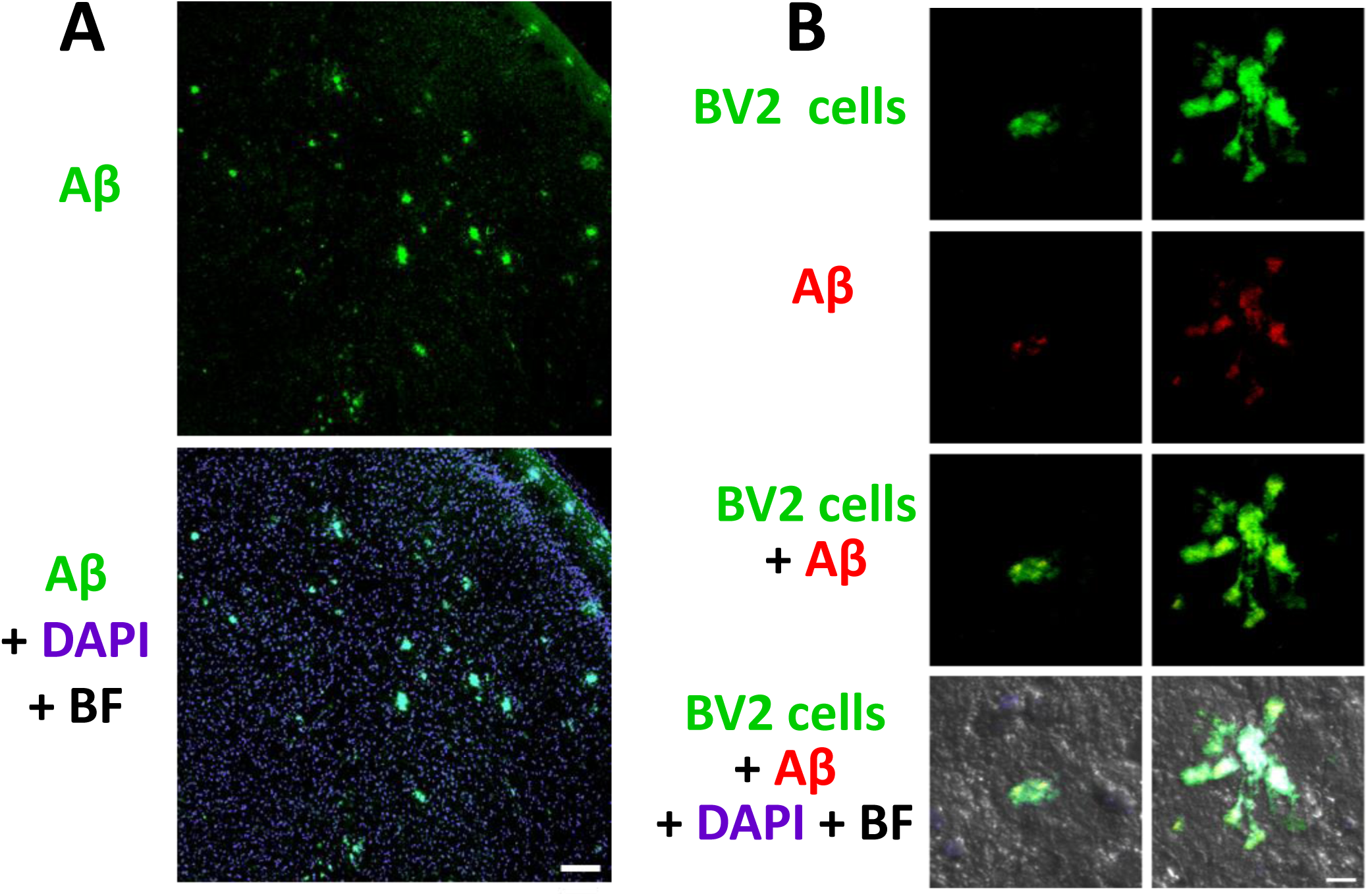
The hybrid protein facilitates higher Aβ uptake and congregation of more GFP-expressing BV2 effector cells around Aβ plaques in APP/PS1 AD mouse brain. (A) Brain sections of APP/PS1 AD mouse at 10 μm thickness was previously verified to have abundant Aβ plaques by Thioflavin (ThT) staining. Scale bar = 100 μm. (B) BV2 cells transfected with GFP plasmid were used as effectors. Phagocytosis assay was carried out by seeding the BV2 effector cells onto a 10 cm plate with AD mouse brain sections mounted on a slide. After three hours incubation at 37°C, the slides were processed for immunohistochemistry and confocal microscopy. Scale bar = 10 μm.

### The hybrid protein facilitates degradation of Aβ by trafficking to the endolysosomal system without compromising cell survival

To distinguish between Aβ uptake and degradation or clearance, we quantified the intracellular Aβ in a time course through confocal microscopy and FACS analysis. Figure 4 shows that BV2 cells effectively clear the phagocytosed Aβ in the presence of the hybrid protein such that a very minimal number of cells associated Aβ are left at the end of 72 hours. The most rapid rate of clearance rate takes place within three hours after Aβ uptake. Trafficking of the internalized Aβ to the phagocytic pathway was evident by its colocalization with the early and late endosomes, and lysosome (Figure 5 for oAβ and Figure 6 for fAβ).

**Fig. 4.**
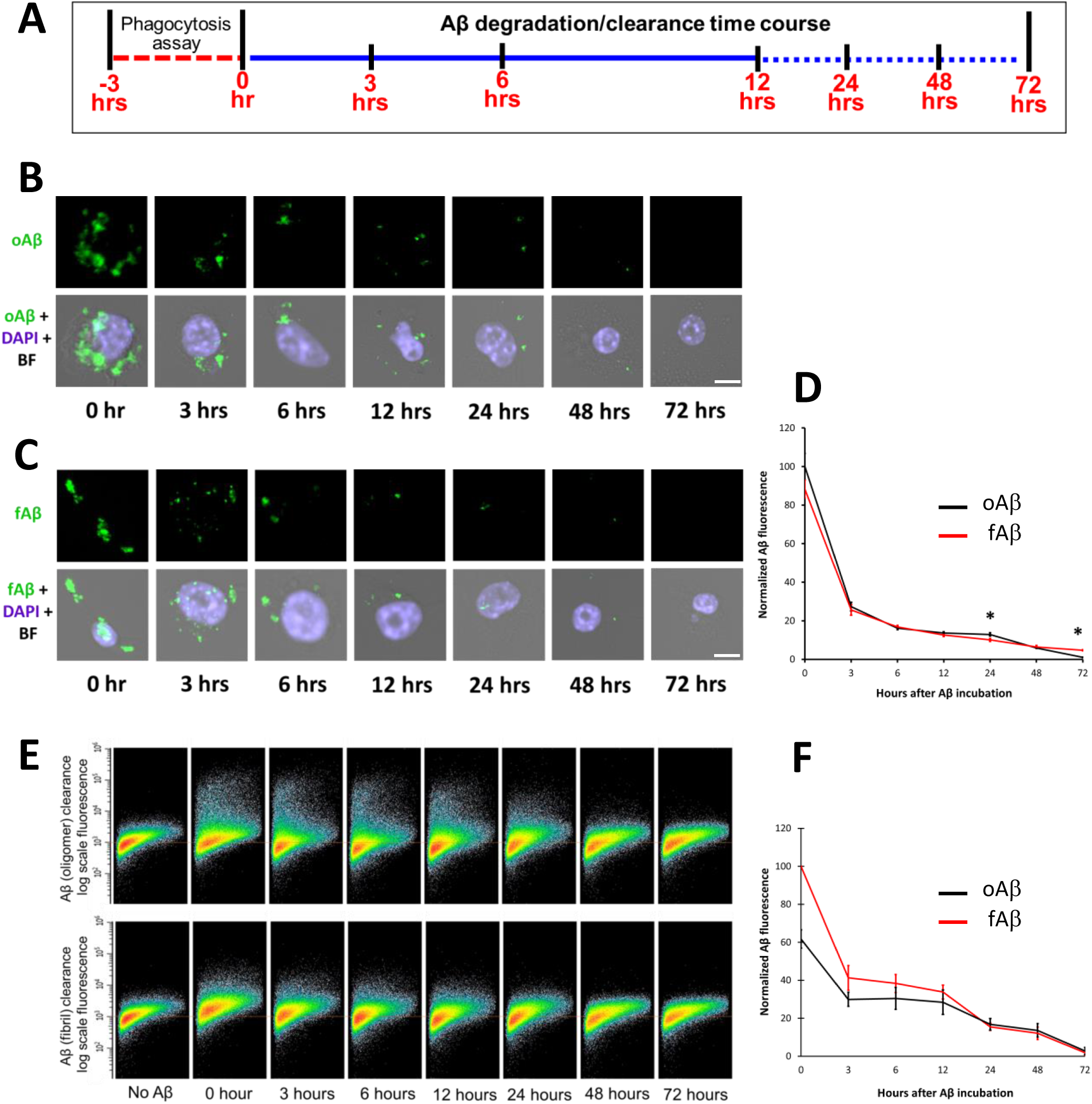
BV2 cells effectively clears phagocytosed Aβ in the presence of the hybrid protein. (A) Schematic diagram of the clearance/degradation assay. BV2 cells were fed with fluorescently-labelled Aβ in the presence of 200nM nAβBPh for three hours followed by trypsinization. The cell associated Aβ at the indicated time points after phagocytosis assay was determined using confocal microscopy and fluorescence-activated cell sorting (FACS) (B-D) Confocal analysis showed progressive reduction of intracellular Aβ as shown in the representative images for oAβ (B) and fAβ (C). (D) Quantification using ImageJ Fiji. Scale bar = 10 μm. n = 18 fields with at least 20 cells/field (source data **Supplementary Figure 4**). FACS analysis showed progressive reduction of cell-associated Aβ as shown in the density plot (E) and quantification (F). The Y-axis is Aβ fluorescence in logarithmic scale. n = 5 independent experiments with at least 30,000 cells each. Asterisk (*) denotes significant differences at P ≤ 0.05 between the means of oAβ and fAβ at the same time point using unpaired, two-tailed Student’s t-test. Data are expressed as mean ± SEM.

**Figure 5.**
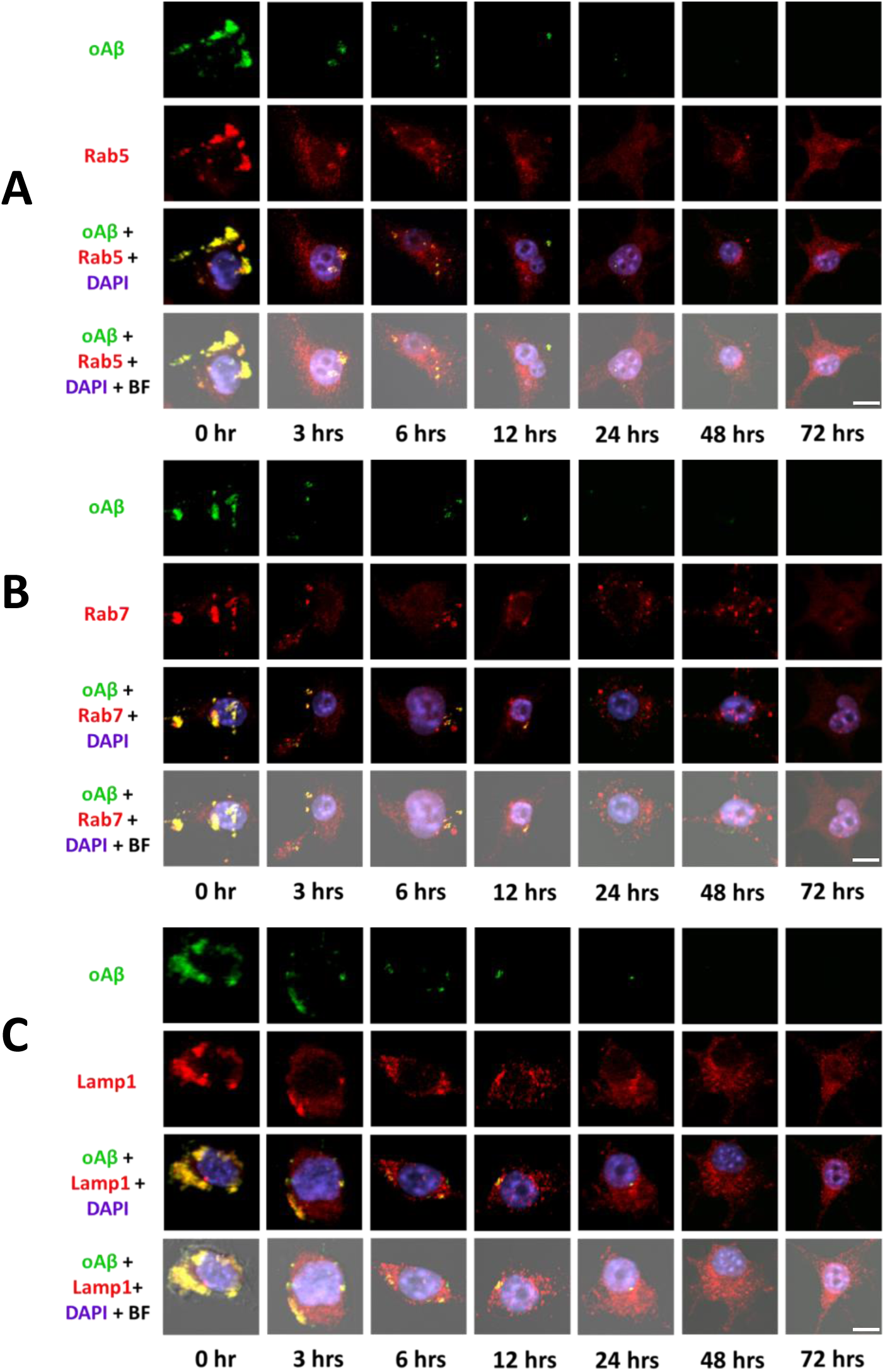
Phagocytosed Aβ oligomers (oAβ) are trafficked to the endolysosomal system for degradation. Representative micrographs show colocalization of oAβ (green) with (A) early endosomal (Rab5), (B) late endosomal (Rab7), and (C) lysosomal (Lamp1) markers (red) in DAPI (blue)-stained BV2 cells following three hours phagocytosis across different time points during degradation. Colocalization in the merged slices appear yellow. Note the high colocalization of Aβ with the early and late endosomes, and the lysosomes, and the progressive decrease in Aβ signal that implies its trafficking to the endolysosomal system and its eventual degradation. Scale bar = 10 μm.

**Figure 6.**
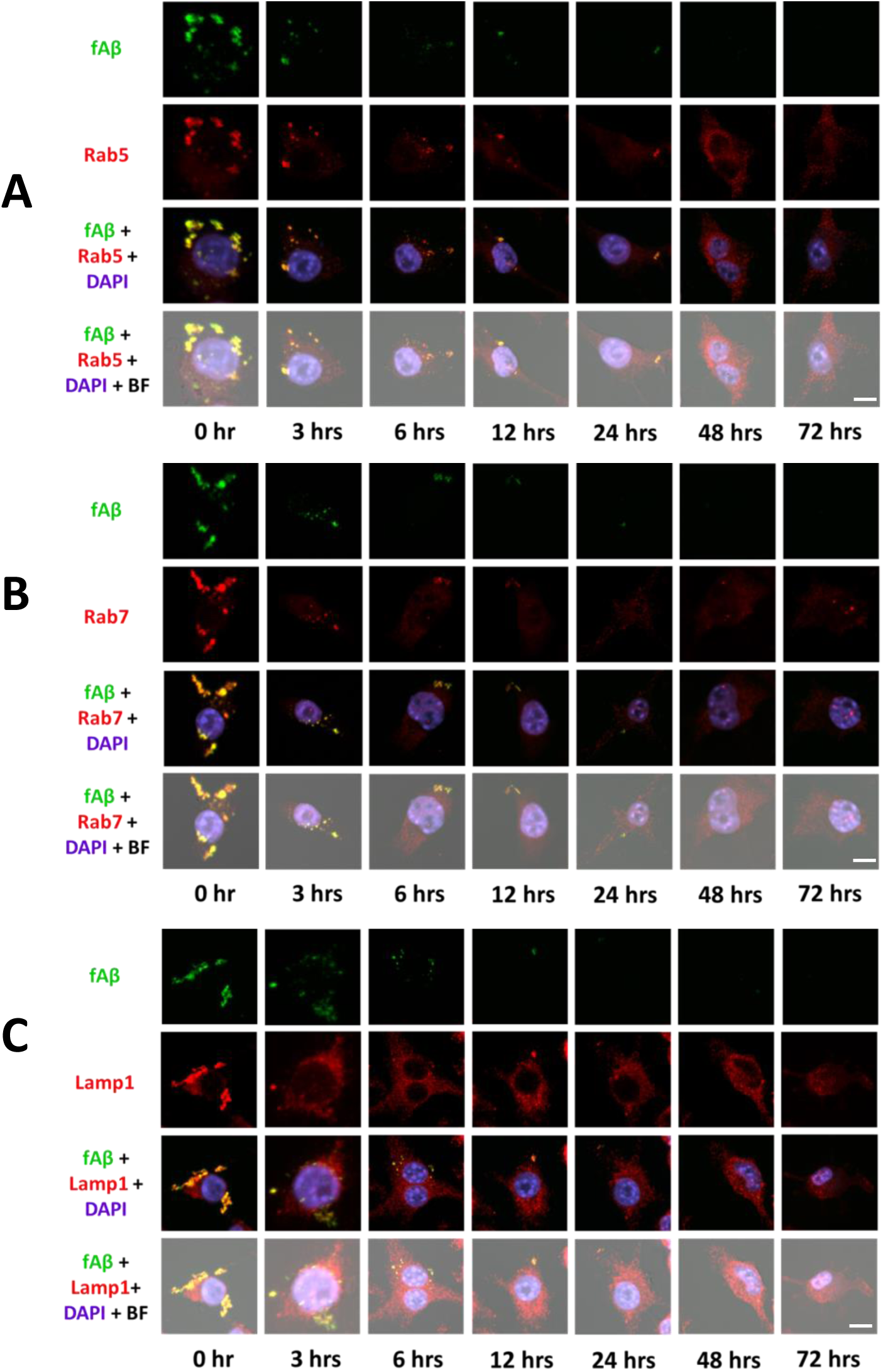
Phagocytosed Aβ fibers (fAβ) are trafficked to the endolysosomal system for degradation. Representative micrographs show colocalization of fAβ (green) with (A) early endosomal (Rab5), (B) late endosomal (Rab7), and (C) lysosomal (Lamp1) markers (red) in DAPI (blue)-stained BV2 cells following three hours phagocytosis across different time points during degradation. Colocalization in the merged slices appear yellow. Note the high colocalization of Aβ with the early and late endosomes, and the lysosomes, and the progressive decrease in Aβ signal that implies its trafficking to the endolysosomal system and its eventual degradation. Scale bar = 10 μm.

### Phagocytic clearance of Aβ has no significant effect on the survival of BV2 microglial cells

Since microglial clearance of Aβ in AD brain eventually leads to massive loss of cells, we also compared the viability of the cells in the presence or absence of the hybrid protein using Water Soluble Tetrazolium (WST)-8-based colorimetric assay. The robust hybrid protein-mediated internalization and eventual degradation of Aβ did not affect the survival of the BV2 cells until 72 hours (Supplementary Fig. 5)

### Aβ uptake facilitated by the hybrid is specific to MerTK receptor

To interrogate the specificity of hybrid protein-mediated phagocytosis of Aβ by BV2 cells through MerTK receptor, we performed competition and blocking studies. The Aβ uptake of BV2 cells in the presence of the hybrid protein is significantly reduced upon the addition of Mer-Fc, the soluble ligand binding domain of MerTK. However, no significant reduction in the uptake of Aβ is observed upon the addition of RAGE-Fc, the soluble ligand binding domain of the RAGE receptor (Figure 7A and B, left).

**Figure 7.**
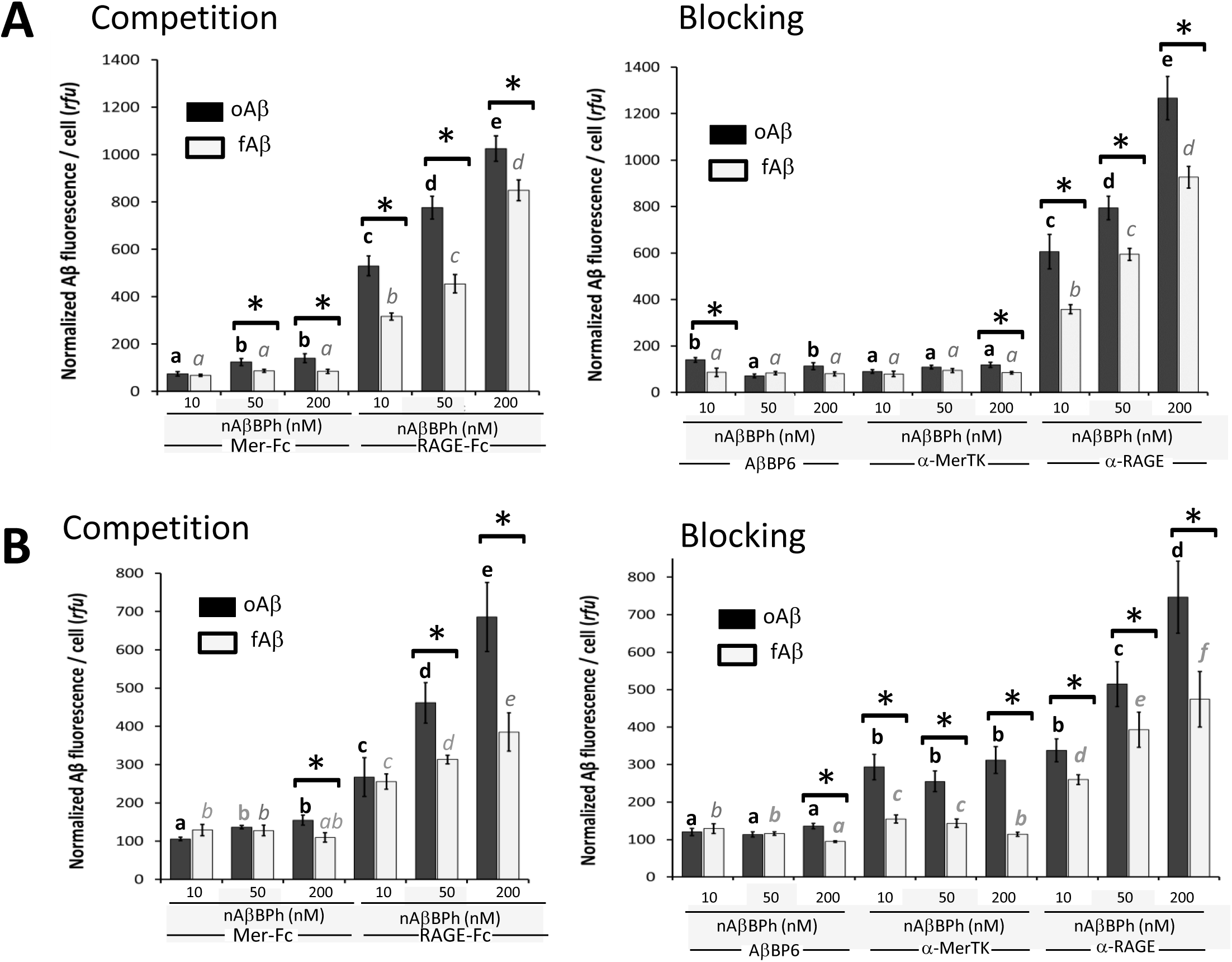
Competition and blocking studies show that Aβ uptake facilitated by the hybrid is specific to MerTK receptor. (A) Confocal and (B) FACS revealed that the fluorescence signal of internalized Aβ was significantly reduced upon the addition of Mer-Fc but not upon the addition of RAGE-Fc (left). A similar reduction was observed after dominant negative blocking through the addition of excessive AβBP6. Blocking with α-MerTK antibody also significantly reduced Aβ internalization, whereas blocking with α-RAGE antibody had no effect on Aβ uptake of BV2 cells in the presence of the hybrid protein (right). Fluorescence data were normalized to the negative control (0 nM nAβBPh) following background subtraction using cells not fed with Aβ. n = 10 fields with at least 20 cells/field for confocal analysis and n = 6 for FACS with at least 30,000 cells. Different letters denote statistical significance at P ≤ 0.05 between treatments using unpaired, two-tailed Student’s t-test. Asterisk (*) denotes significant differences between the means of oAβ and fAβ in the same treatment. Data are expressed as mean ± SEM.

A similar reduction to MerFc competition is observed when dominant negative blocking is done by adding excessive amount of AβBP that binds Aβ but has no MPD for MerTK recognition. Further, blocking using α-MerTK antibodies that bind to the extracellular domain of the receptor also significantly reduced the amount of Aβ internalized by BV2 cells. This reduction in cell associated Aβ was not observed upon blocking with the use of α-RAGE antibodies that bind to the extracellular domain of RAGE (Figure 7A and B, right).

To visualize the binding of the hybrid protein to the MerTK receptor at the cell surface, surface binding assay was done by performing phagocytosis at 4°C. The 4°C incubation temperature allows the interaction of the hybrid protein with both Aβ and MerTK but prevents the internalization of the Aβ cargo even if the ligand is bound to the receptor. The presence of the hybrid protein results to a significant increase in Aβ fluorescence colocalizing with MerTK at the surface of BV2 cells in a concentration dependent manner (Figure 8A-B).

**Figure 8.**
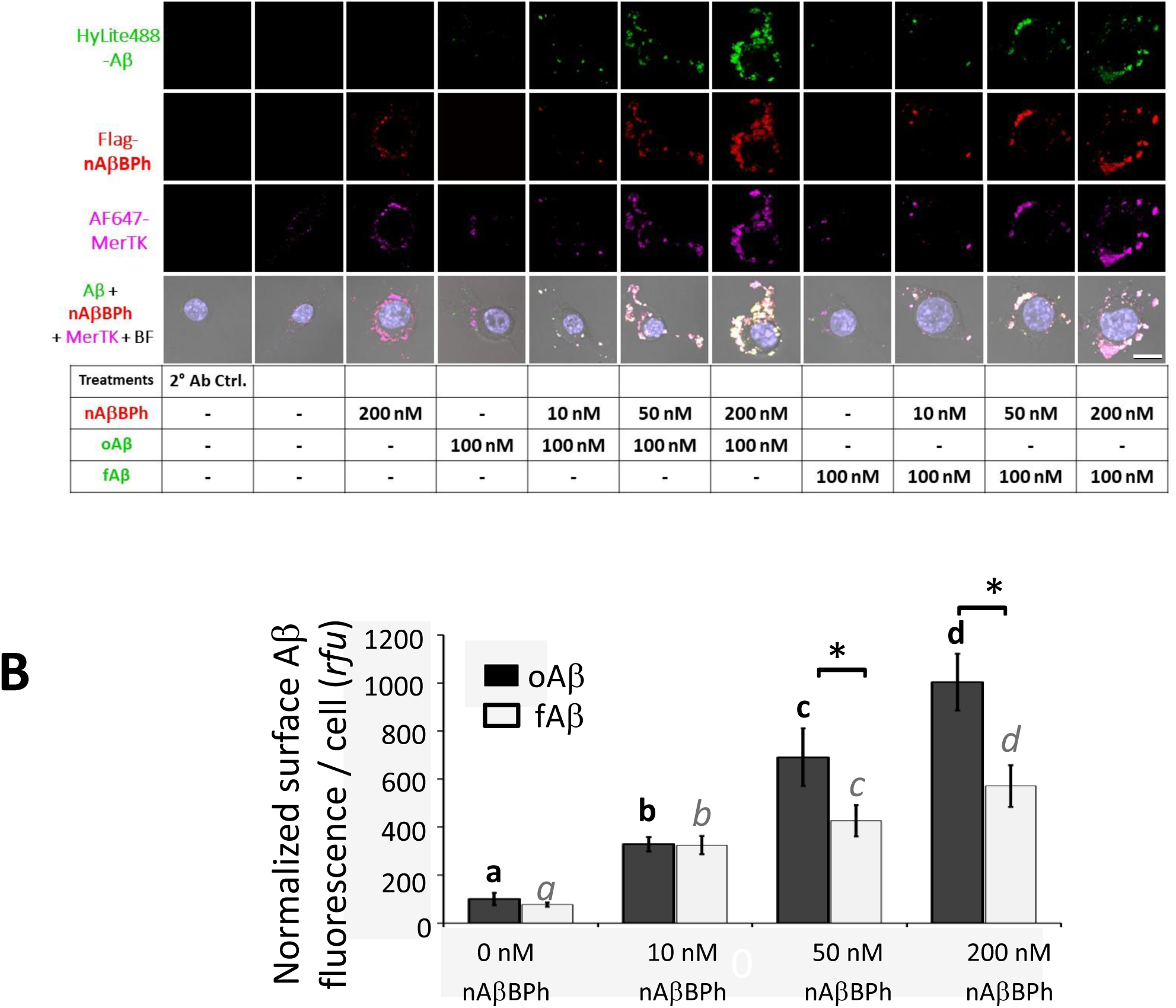
As hybrid concentration increases, the fluorescence of Aβ colocalizing with MerTK receptor at the surface of BV2 cells also increases. (A) Representative micrographs and (B) quantification showed the colocalization. Normalized surface Aβ fluorescence / cell (*rfu*) of FLAG-tagged hybrid protein (red) with bound Aβ (green) and MerTK receptor (pink). BV2 cells were incubated with HiLyte488 Aβ in the presence or absence of FLAG-tagged nAβBPh at 4°C for 3 hours. After the cells were washed, FLAG-nAβBPh was detected by immunocytochemistry using anti-FLAG primary antibodies (Abs) and AlexaFluor 568 secondary Abs and MerTK was detected using anti-MerTK Abs conjugated to Alexa Fluor 647. Fluorescence data were normalized to the negative control for oAβ (0 nM nAβBPh) following background subtraction using cells not fed with Aβ. Different letters denote statistical significance at P ≤ 0.05 between treatments using unpaired, two-tailed Student’s t-test. Asterisk (*) denotes significant differences between the means of oAβ and fAβ in the same treatment. Data are expressed as mean ± SEM. Scale bar = 5 μm.

Since MerTK activation is characterized by the rearrangement of the cytoskeleton leading to the redistribution and colocalization of non-muscle myosin IIA (NMMIIA) with the ingested cargo (Caberoy et al., 2010a; Caberoy et al., 2012a; Strick et al., 2009), we looked at the effect of the hybrid protein on NMMIIA rearrangement and colocalization with Aβ. Figure 9 shows that the presence of the hybrid protein and of the N-terminal of Tubby with the MPD that recognizes MerTK results to the rearrangement of NMMIIA, but not the presence of AβBP and Aβ only.

**Figure 9.**
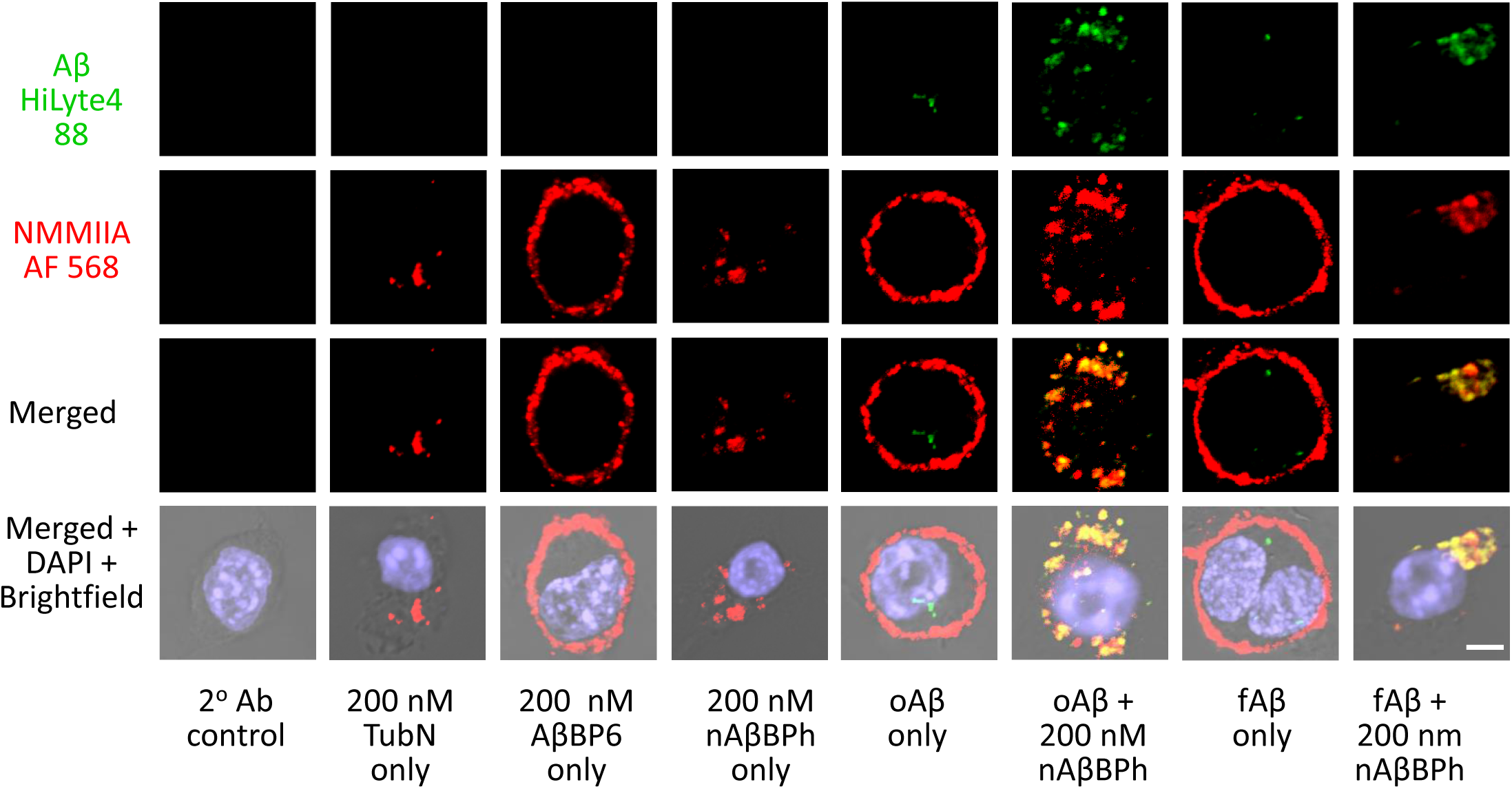
The hybrid-mediated phagocytosis of Aβ results to cytoskeletal rearrangement and its colocalization with non-muscle myosin IIA (NMMIIA). Representative micrographs showing the rearrangement of non-muscle myosin IIA in the presence of the hybrid protein and its colocalization with the internalized Aβ. BV2 cells were incubated with HiLyte488 Aβ in the presence or absence of the hybrid protein at 37°C for 3 hours. After the cells were washed, NMMIIA was detected by immunocytochemistry using anti-NMMIIA primary Abs and AlexaFluor 568 secondary Abs. Scale bar = 5 μm.

### The hybrid protein significantly reduces the production of TNF-α, IL-1β in BV2 cells after Aβ stimulation

Since the expression of pro-inflammatory cytokines is induced by Aβ in activated microglial cells (Bertram and Tanzi, 2019; Wolfe, 2016; Zenaro et al., 2017) we would like to find out whether the hybrid protein that effectively facilitated Aβ phagocytosis through MerTK can reduce the production of inflammatory and oxidative factors in BV2 cells. First, we quantified the protein expression of selected pro- and anti-inflammatory cytokines by multiplex ELISA. Addition of the hybrid protein to BV2 cells incubated with 1 *µ*M Aβ for 24 hours resulted to a significant reduction in the protein expression levels of TNF-α, IL-1β and IL-10 (Figure 10A). This significant reduction in secreted TNF-α is consistent with the intracellular TNF-α in BV2 cells after six hours incubation with 1 *µ*M oAβ (Figure 10B).

**Figure 10.**
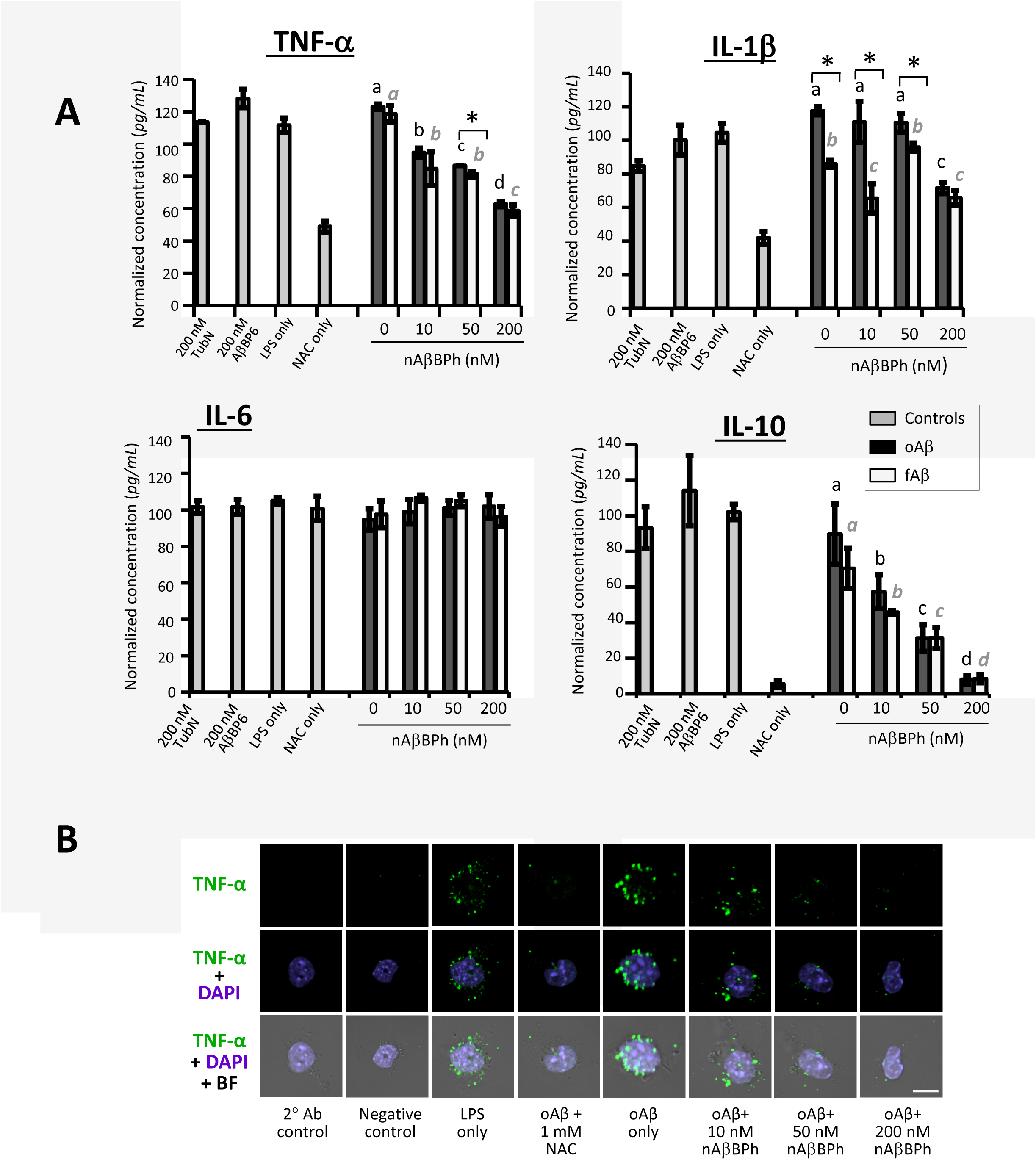
The hybrid protein significantly reduces the production of TNF-α, IL-1β, and IL-10 in BV2 cells after Aβ stimulation. (A) TNF-α, IL-1β, IL-6, and IL-10 in the cell culture media of BV2 cells treated with 1µM Aβ oligomer (oAβ) or Aβ fibril (fAβ) for 24 hours in the presence of nAβBPh was quantified by multiplex ELISA using the mouse high sensitivity T-Cell discover array 18-plex (MDHSTC18) of EveTechnologies. LPS and N-acetyl cysteine were used as reference. TubN and AβBP6, proteins that contain part of the hybrid, were likewise included. n = 3 technical replicates. Different letters denote statistical significance at P ≤ 0.05 between treatments using two-tailed Student’s t-test. Asterisk (*) denotes significant differences between the means of oAβ and fAβ in the same treatment. Data are expressed as mean ± SEM. (B) Representative confocal images show that the hybrid protein significantly reduces intracellular TNF-α in BV2 cells after oAβ stimulation. Scale bar = 5 *µ*m.

### Nitrite and ROS production of BV2 cells in response to Aβ stimulation is significantly reduced in the presence of the hybrid protein

An increased inducible nitric oxide synthase (iNOS) expression leading to elevated levels of nitric oxide (NO) in glial cells during Aβ elicited inflammatory and immune responses is well documented. So, we next determined the concentration of NO through the formation of its stable metabolite nitrite using Greiss assay. Nitrite accumulation, hence, NO production, was significantly reduced in a concentration-dependent manner when BV2 cells were treated with Aβ to induce a proinflammatory response for 24 hours (Figure 11A).

**Figure 11.**
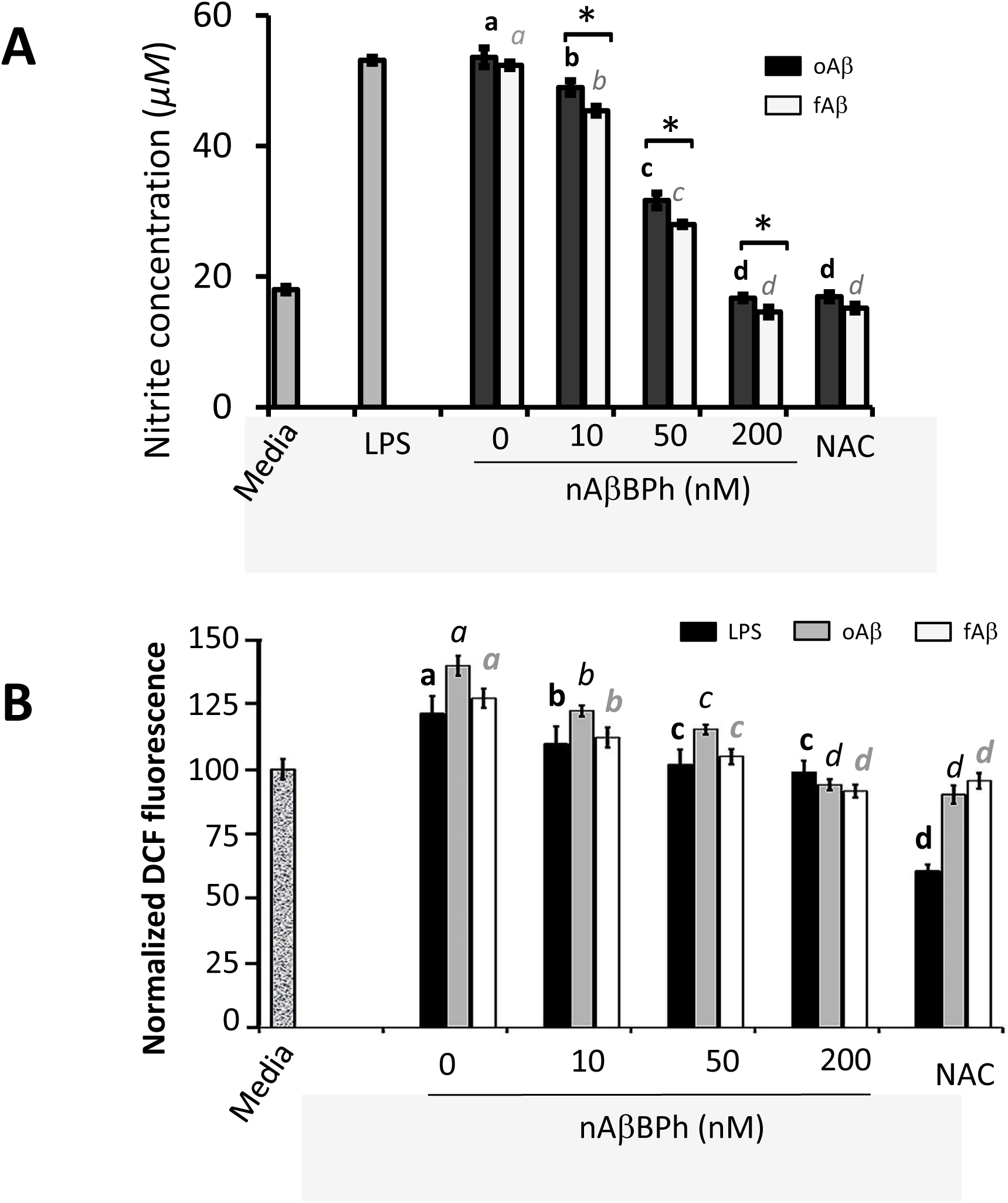
The hybrid protein significantly reduces nitrite and reactive oxygen species (ROS) productions of BV2 cells in response to Aβ stimulation. (A) Nitrite concentration in the media was quantified using Greiss assay. Lipopolysaccharide (LPS: 1µg/mL) and N-acetyl cysteine (NAC: 1mM) were used as positive and negative controls, respectively. (B) ROS assay. BV2 cells were treated with 1µM Aβ oligomer (oAβ) or Aβ fibril (fAβ) in the presence of the hybrid protein. DCF fluorescence was quantified using fluorescence plate reader at 485/535 nm excitation/emission wavelengths. LPS was used as control and N-acetyl cysteine, a known antioxidant, was used as reference. n = 6 technical replicates, a representative of 3 experimental replicates. Different letters denote statistical significance at P ≤ 0.05 between treatments using two-tailed Student’s t-test. Asterisk (*) denotes significant differences between the means of oAβ and fAβ in the same treatment. Data are expressed as mean ± SEM.

Lastly, we quantified the level of cellular reactive oxygen species (ROS) using 2’,7’- dichlorofluorescin diacetate (DCFDA) (Giovanna et al., 2010; Schilling and Eder, 2011), a cell permeant dye that measures hydroxyl, peroxyl and other ROS activity within the cell through its subsequent oxidation by ROS into a highly fluorescent compound dichlorofluorescin (DCF). ROS based on DCF fluorescence intensity of BV2 cells treated with 1 *µ*M Aβ for 24 hours was also significantly reduced in the presence of the hybrid protein in a concentration-dependent manner (Figure 11B). These observations suggest that the presence of the hybrid protein significantly reduces the levels of Aβ-induced expressions of inflammatory and oxidative factors in BV2 cells.

### The hybrid protein crosses the blood brain barrier (BBB)

Since brain is the target site for treatment, we tested whether the hybrid protein could cross the BBB. Thirty minutes after injection, we detected the FLAG-tagged hybrid protein by Western blot in the total lysate of the right hemisphere of the brain of mice injected intraperitoneally with the hybrid protein but not in the control mice injected with PBS (Figure 12A). To validate that the hybrid protein had really entered the brain tissues and was not just in the blood vessels within the brain, we performed immunohistochemistry (IHC) on the left hemisphere of the brain and examined the presence of the hybrid protein in the hippocampus. The representative confocal images in Figure 12B and C show that the hybrid protein colocalizes with both the microglia and MerTK receptor in the hippocampus, confirming that it has crossed the BBB.

**Figure 12.**
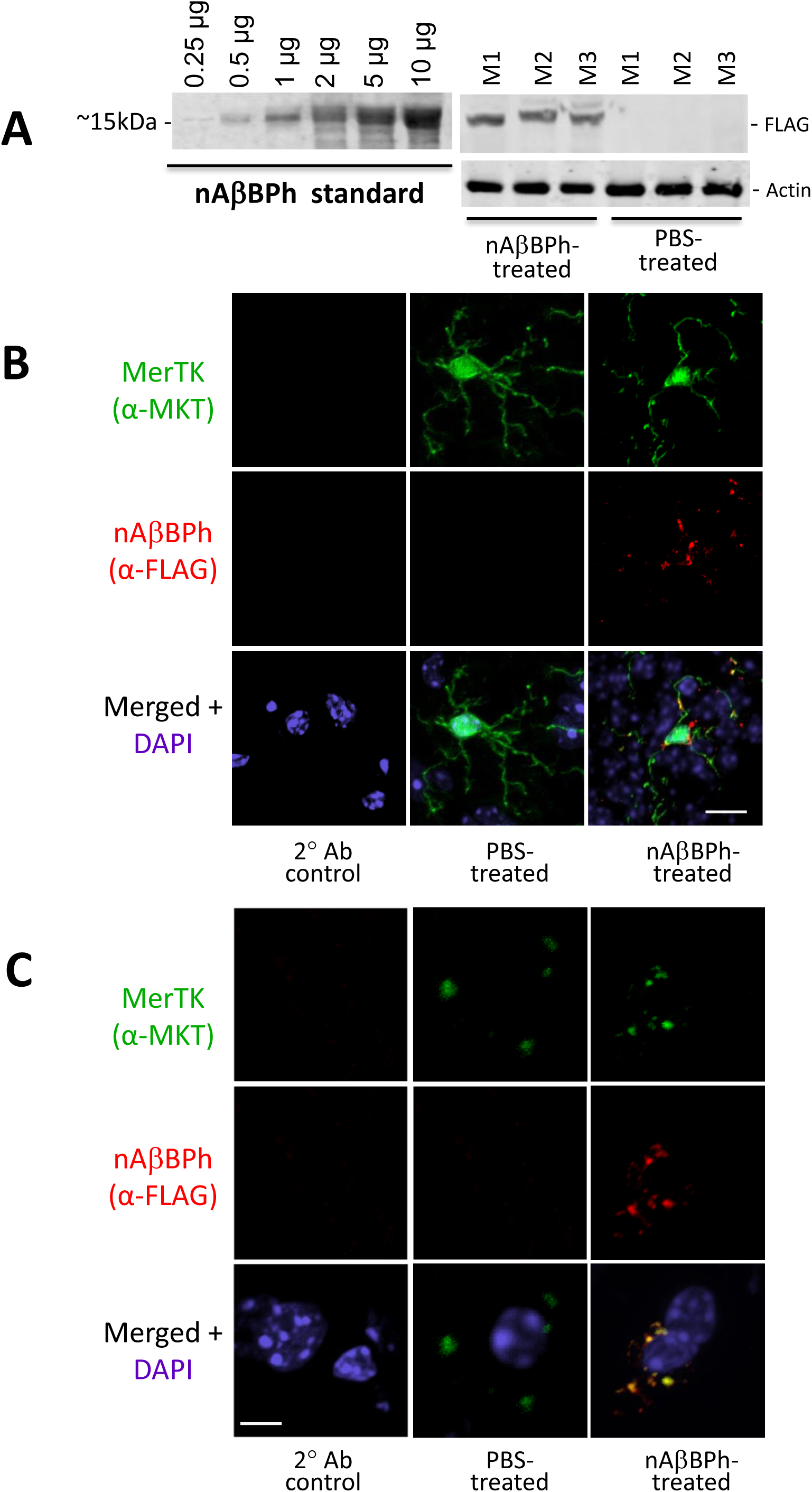
The hybrid protein crosses the blood brain barrier (BBB) 30 minutes after intraperitoneal injection. (A) Western blot analysis detects the FLAG-tagged hybrid protein in the whole brain lysate of hybrid-treated mice but not in PBS-treated mice. (B) The hybrid colocalizes with the microglia and with MerTK receptor (C) in the hippocampus. Colocalization in the merged slices appear yellow. Scale bar = 5 *µ*m in (B) and 2 *µ*m in (C).

### The hybrid protein causes a significant reduction in the Aβ burden in the brain of female APP/PS1 AD mice after 2 months of treatment

Since the hybrid protein can cross the BBB, the next thing we wanted to know is whether hybrid protein treatment can reduce the level of Aβ in the brain of AD mice. We used an APP/PS1 humanized mouse model of AD. This humanized AD mouse is homozygous for two mutant alleles of human Amyloid Precursor Protein Swedish (APP Swe) mutation and Presenilin 1 (Psen1 that codes for an integral component of γ-secretase enzyme complex that cleaves APP). These mutations in APP and Psen1 result to the manifestation of characteristic Aβ plaque pathology as early as 6-7 months (Jankowsky et al., 2004). Since at around six months this AD mouse model already has Aβ plaques in the brain, we used 6-months old mice for treatment.

We quantified the levels of Aβ 40 and 42 in the brain by sandwich ELISA since Aβ40 and Aβ42 are the most abundant species of Aβ found in plaques in AD brain (Murphy and LeVine, 2010; Selkoe, 2001). Figure 13A shows a significant reduction in the levels of insoluble and total Aβ42 as well as the total Aβ42/Aβ40 ratio in the brain of female mice after 60 days of hybrid protein treatment. Complementarily, this reduction in Aβ load in the brain of female mice is also observed using IHC as shown in Fig. 13b where relatively fewer and smaller Aβ plaques are detected in both the cortex and the hippocampus.

**Figure 13.**
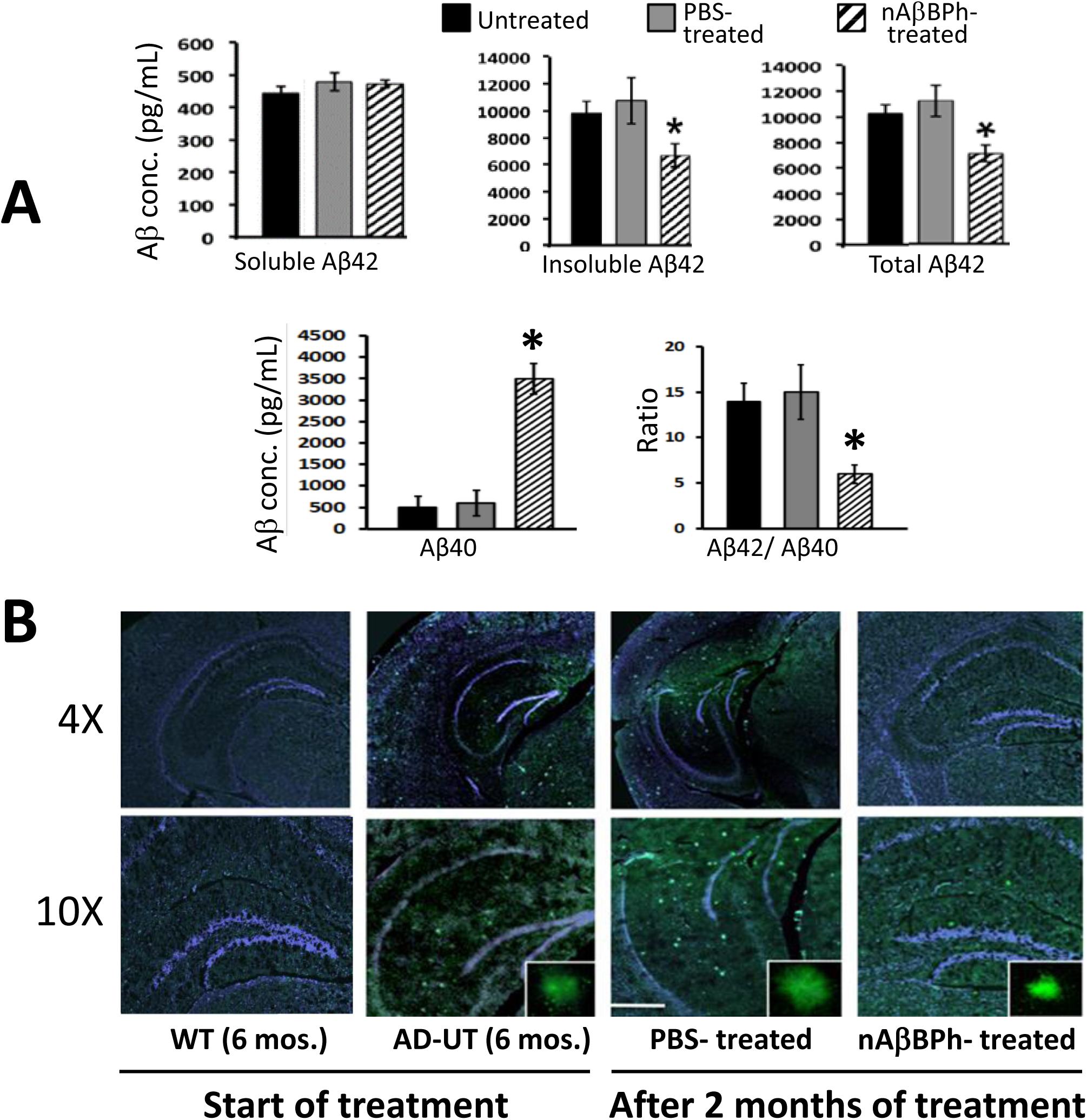
Hybrid treatment for 2 months reduces Aβ burden in the brain of APP/PS1 AD mice. (A) Insoluble and total Aβ42, and the ratio of total Aβ42/Aβ40 are significantly reduced in the brain of female mice based on sandwich ELISA using Aβ40 and Aβ42 EZ brain ELISA Kit (EMD Millipore) following the manufacturer’s protocol. n = 6 mice. Asterisk (*) denotes significant difference at P ≤ 0.05 using two-tailed Student’s t-test. Data are expressed as mean ± SEM. (B) Representative micrographs show marked reduction in the size and number of Aβ plaques detected through thioflavin (ThT) staining in of the brain of female mice injected with the hybrid compared to the control. AD-UT = untreated APP/PS1. Scale bar = 50 *µ*m.

### Hybrid protein treatment reduced IL-6 level in the brain of AD mice

Aβ plays a pivotal role in the progression of Alzheimer’s disease through its neurotoxic and inflammatory effects (Yu and Ye, 2015). Microglial cells activated by Aβ induce the expression of pro-inflammatory cytokines including interleukin (IL)-1β, IL-6, IL-8, tumor necrosis factor-α (TNF-α), chemokines and reactive oxygen and nitrogen species (Wolfe, 2016). Hence, we quantified the concentration of representative cytokines in the brain in order to assess whether the reduction in Aβ burden in the brain as a result of hybrid protein treatment also translates to a reduction in the levels of proinflammatory cytokines. Fig.14 shows that hybrid protein administration for two months results to a reduction in the levels of IL-6, while the levels of TNF-α, IL-1β, and IL-10 did not change significantly.

**Figure 14.**
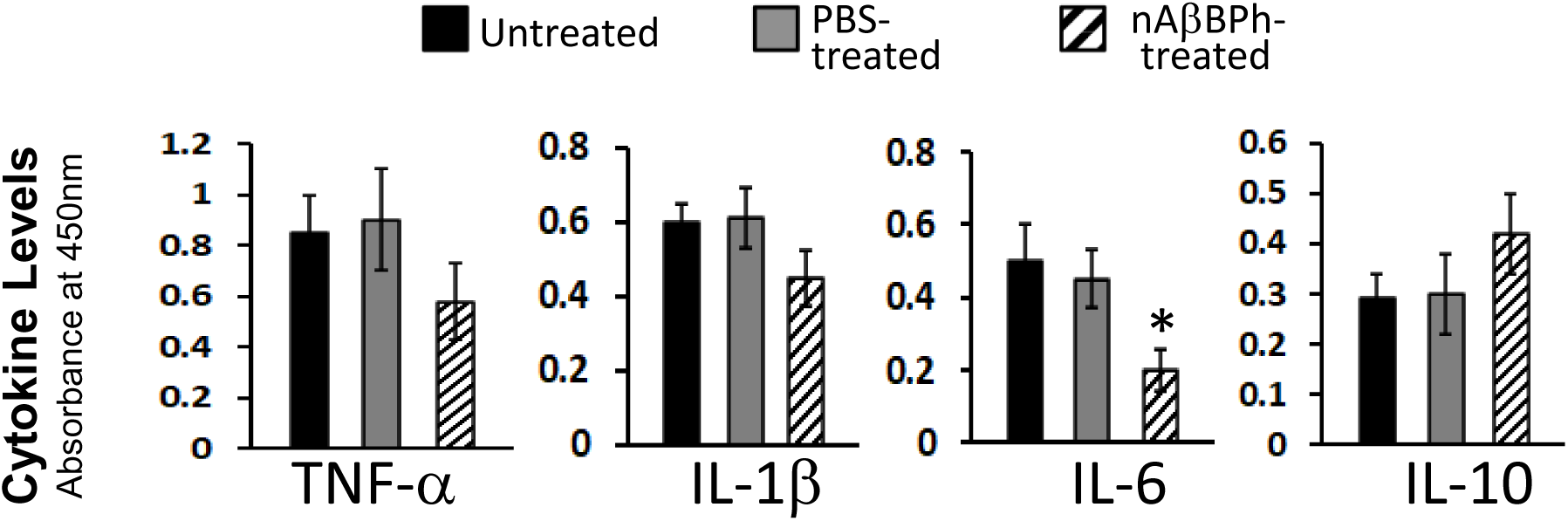
Hybrid treatment for 2 months reduces the level of IL-6 in the brain of APP/PS1 AD mice. ELISA was done using the Qiagen Multi-Analyte ELISArray kit following the manufacturer’s protocol to quantify the protein expression levels of representative cytokines in the brain homogenates. n = 6 mice. Asterisk (*) denotes significant difference at P ≤ 0.05 using two-tailed Student’s t-test. Data are expressed as mean ± SEM.

### Treatment of the hybrid protein protects the nerve cells from dying

Since treatment of the hybrid protein results to a decrease of both Aβ burden in the brain and IL-6 cytokine whose expression is upregulated in AD mice and AD patients, we wanted to know whether these effects result to the protection of the brain cells from dying. So, we performed IHC to detect the different cell types in the hippocampus using specific markers. These include MAP2 (an isoform of microtubule specific protein expressed only in the neuron), IBA1 or ionized calcium binding adaptor molecule 1 (a microglial and macrophage-specific calcium-binding protein that is involved with phagocytosis) and GFAP or glial fibrillary acidic protein (an intermediate filament protein that is expressed by astrocytes in the central nervous system). Figure 15 shows the brain cell profile in the hippocampus of AD mice. After two months treatment, we see that more nerve cells are still present in the brain of mice injected with the hybrid protein compared to the control injected only with PBS where we see a dramatic reduction of neurons. This result implies that the hybrid protein protects the nerve cells from dying.

**Figure 15.**
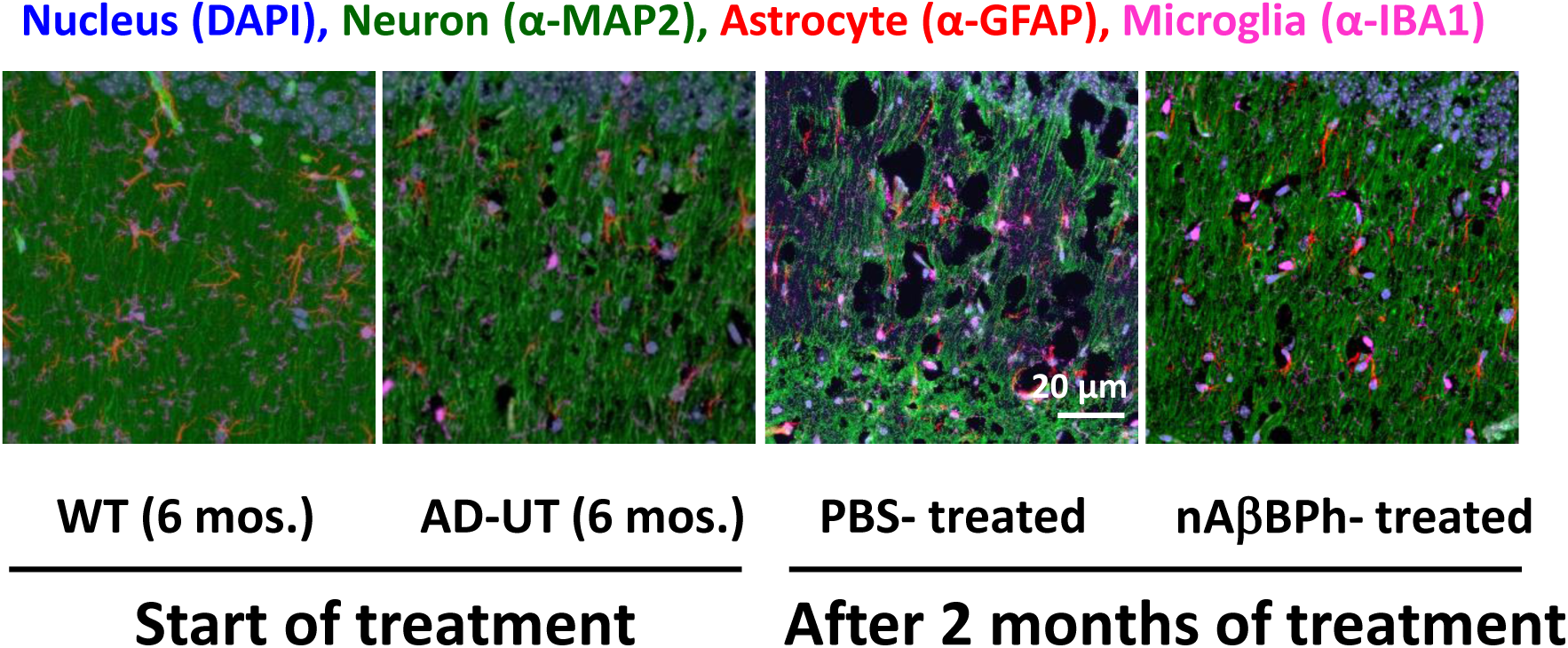
Hybrid treatment for 2 months protects the nerve cells from dying. Representative micrographs of the hippocampus region of APP/PS1 AD mice show that after 2 months treatment, more nerve cells are still present in the brain of hybrid-treated mice compared to the PBS-treated control in which a marked reduction in the number of nerve cells is evident. Brain sections were processed by immunohistochemistry and brain cells were detected under the confocal microscope using the following cell specific antibodies with the corresponding secondary antibodies: neurons (α-MAP2 and AF488), microglia (α-IBA1 and AF 657), and astrocytes (α-GFAP and AF568). AD-UT = untreated APP/PS1. n = 6 mice.

### Hybrid protein treatment does not cause histopathological toxicity to AD mice

Aside from mortality, liver toxicity and kidney injury following drug treatments are the first indicators of drug safety (Faria et al., 2019; Liu et al., 2011). To assess the safety or toxicity of the hybrid protein, we performed thorough histological analysis of the liver and kidney using Hematoxylin and Eosin (H and E) staining, the most commonly used method in medical diagnosis and considered the gold standard in histopathological analysis. We did not see histological abnormalities such as infiltration of immune cells, inflammation, fibrosis, and degeneration or necrosis that are indications of hepatic and renal toxicities (Supplementary Figure 6). So aside from 100% survival of all hybrid protein-treated mice, the normal histological features of both the liver and kidney of treated mice suggests that the hybrid protein is safe and is well-tolerated by APP/PS1 AD mice even if administered daily at 50 mg/kg body weight for two months.

## Discussion

The accumulation Aβ peptides within the brain arises from an imbalance of the production and clearance of Aβ. Instead of the increased production of Aβ, recent evidence indicates that in most cases of AD, Aβ accumulation results because of an impaired clearance mechanism (Kurz and Perneczky, 2011; Mawuenyega et al., 2010; Ramanathan et al., 2015; Wildsmith et al., 2013). Hence, methods that modulate Aβ clearance have been important targets in the development of disease modifying therapeutic agents for Alzheimer’s disease (Mawuenyega et al, 2010).

Microglial clearance is one of the critical pathways to remove the Aβ in AD (Hickman et al., 2008; He et al., 2020; Zuroff et al., 2017). Oligomeric and fibrillar Aβ are primarily internalized though receptor-mediated endocytosis, via a number of receptors that are expressed on the surface of microglia and astrocytes (Ries and Sastre, 2016).

Here, we have shown that the hybrid protein mediated the clearance of Aβ by BV2 cells through MerTK receptor. The presence of the hybrid protein significantly improved the phagocytosis efficiency of both oligomeric and fibrillar Aβ by BV2 cells in a concentration-dependent manner. The presence of the hybrid protein resulted not only to a significantly higher level of Aβ uptake but also to a significantly higher percentage of phagocytic cells. The robust Aβ phagocytosis facilitated by the hybrid protein is consistently better for Aβ oligomers than fibrils, considering that the Aβ binding portion of the hybrid protein was chosen because of its highest affinity to oAβ. This result is very promising since the soluble oligomeric Aβ assemblies are more neurotoxic and are implicated in causing synaptic dysfunction, neurodegeneration and cell death than the fibrillar form (Pan et al., 2011; Nirmalraj et al., 2020; Koffie et al., 2009; Lacor et al., 2007; Lambert et al., 1998; Pickett et al., 2016; Querol-Vilaseca et al., 2019; Shankar et al., 2008; Walsh et al., 2002; Wang et al., 2002).

In the absence of the hybrid protein, the baseline phagocytic function of BV2 cells is not significantly different between oAβ and fAβ. Previously, Pan et al. (2011) noted that exposure of BV2 cells for 30 minutes to fAβ resulted in significantly higher microsphere uptake and percentage of phagocytic cells compared to oAβ which led them to suggest that oAβ impairs the microglial phagocytic function. Our results indicate that the hybrid protein does not only facilitate better phagocytic efficiency in BV2 cells but it also helps overcome the effects of oAβ that compromises the cell’s ability for phagocytosis. These findings have significant implications on microglial dysregulation by oAβ that causes proinflammatory and oxidative stress thereby preventing clearance of fAβ deposits, leading to an initial neurodegenerative process characteristic of AD.

Both Aβ oligomers and fibrils phagocytosed in the presence of the hybrid protein are effectively cleared through the endolysosomal system within 72 hours. The fastest rate of clearance takes place in the first three hours after Aβ uptake. Phagocytosed Aβ is trafficked to the early endosome and late endosome, and eventually degraded in the lysosome. The lysosome is the endpoint for endosomes formed by receptor-mediated endocytosis. The lysosome contains hydrolytic enzymes capable of degrading Aβ; however, this machinery is often overwhelmed in AD as the Aβ load exceeds the organelle’s degradation capacity (Zuroff et al., 2017; Gowrishankar et al., 2015; Umeda et al., 2011; Yang et al., 1998).

While the hybrid protein facilitated robust Aβ uptake, we also demonstrated that the high level of ingested Aβ did not result to cell mortality. We have shown in our time course experiments that cell survival of BV2 cells was not significantly reduced up to 72 hours post phagocytosis assay. This result is consistent with the observation that oAβ and fAβ are cytotoxic only at concentrations ≥ 5 µM and ≥ 10 µM, respectively for both BV2 and primary microglia (Pan et al., 2011). However, the decreasing trend in the survival of BV2 cells treated by oAβ, though not statistically significant until 72 hours, may indicate that the cells are starting to be compromised. Interrogation of cell viability over a longer duration might reveal significant cell mortality as Aβ toxicity resulting to cell death progresses slowly.

The hybrid protein offers hope for AD treatment through Aβ clearance because aside from promoting a significantly higher phagocytic efficiency in BV2 using synthetic Aβ, it also facilitates uptake of physiologically generated Aβ. In fact, our functional assay shows that BV2 effector cells in the presence of the hybrid protein are seen to cluster around large plaques in the brain of APP/PS1 AD mouse. The same observation has been noted in AD mouse brain where Aβ plaques are surrounded by microglia which play an important role in limiting the growth of Aβ plaques and plaque-associated disruption of neuronal connection (Bolmont et al., 2008; Meyer-Luehmann et al., 2008; Zhao et al., 2017), and also in the post-mortem brain of AD patients where activated microglia are associated with neuritic plaques (Zuroff et al., 2017; Haga et al., 1989; Mattiace et al., 1990; Rozemuller et al., 1986). The induced cellular phagocytosis of Aβ deposits in the brain when co-cultured with effector cells was likewise observed in an amyloid-targeting monoclonal antibody, Gantenerumab (Bohrmann et al., 2012), which is now in a stage three clinical trial.

We have also demonstrated that the hybrid protein-facilitated phagocytosis of Aβ is specific to MerTK receptor. Addition of Mer-Fc, the soluble extracellular ligand-binding domain of MerTK, effectively reduced Aβ uptake, whereas addition of RAGE-Fc, the soluble extracellular ligand-binding domain of RAGE, did not have any effect on Aβ uptake of BV2 cells. Similarly, blocking the interaction of the hybrid protein with MerTK using α-MerTK antibodies also reduced the Aβ uptake of BV2 cells, which was not observed when α-RAGE antibodies were used. Additionally, dominant negative blocking by using excessive AβBP that binds Aβ but has no MPD to recognize MerTK also resulted to a significant decrease in Aβ phagocytosis. These results support the effectiveness of the hybrid protein in diverting the phagocytosis of Aβ to MerTK from RAGE, otherwise, the use of a competitor molecule or antibodies to block RAGE should have significantly decreased the uptake of Aβ as observed in previous studies upon blocking of RAGE (Deane et al., 2012). The results of these competition and blocking studies are corroborated by the actual binding of the ligand colocalizing with Aβ to the MerTK receptor at the surface of the cell. Moreover, the colocalization of Aβ with NMMIIA also confirms activation of MerTK receptor. Ligand-induced activation of MerTK results to receptor autophosphorylation followed by the rearrangement of the cytoskeletal proteins and the extension of the plasma membrane for engulfment, leading to the redistribution and colocalization of NMMIIA with the ingested cargo (Caberoy et al., 2010a; Caberoy et al., 2012a; Strick et al., 2009). All these observations suggest that the hybrid protein-facilitated phagocytosis of Aβ is mediated specifically by MerTK receptor.

Since the binding of Aβ to plasma membranes appears to be a promising point of intervention in the events leading to the development of AD (Verdier and Penke, 2004), the fact that the hybrid protein diverts the clearance of Aβ from RAGE and other PRRs whose activation causes the release of inflammatory factors is an additional advantage. The fact that the hybrid protein promotes significantly better uptake of oAβ offers hope for *in vivo* studies in terms of sequestering the soluble and more neurotoxic oAβ first, so that the fAβ that mainly makes up the plaques can be effectively cleared once the oAβ that attenuates microglial phagocytosis of fAβ (Pan et al., 2011) has been dealt with.

The binding of Aβ peptide to cell surface receptors induces proinflammatory gene expression and subsequently cytokine production (Pan et al., 2011). Here, we have shown that the presence of the hybrid protein that facilitated Aβ uptake through the MerTK receptor at the cell surface resulted to a significant reduction of the levels of the proinflammatory cytokines TNF-α and IL-1β. The same reduction was noted in cells treated with N-acetyl cysteine (NAC) known for its antioxidant and anti-inflammatory activities (Cazzola et al., 2017; Faghfouri et al., 2020; Palacio et al., 2011; Sadowska et al., 2007; Uraz et al., 2013; Zhang et al., 2018).

The levels of IL-6 however did not significantly change across different treatments. BV2 cells’ inflammatory activation in response to Aβ stimulation that had been documented thus far included only elevated TNF-α and IL-1β but not IL-6 (Pan et al., 2011; Stansley et al., 2012; Timmerman et al., 2018). It can also be noted that the effect of the hybrid protein in reducing TNF-α after 24 hours culture is more pronounced than IL-1β. Being an immediate response molecule, the maximal response for IL-1β is at three hours whereas the maximal response of TNF-α (and also nitrite) in response to Aβ is at 24 hours (Pan et al., 2011; Kim et al., 2006). This fact may also explain the pronounced difference in intracellular TNF-α detection in the different treatments, but not in the other cytokines.

When we looked at the effect of the hybrid protein in the production of IL-10, a representative anti-inflammatory cytokine, we see that it is also significantly reduced in the presence of the hybrid protein and NAC, having similar pattern as TNF-α and IL-1β. The same NAC-induced suppression of IL-10 secretion has been observed in LPS-challenged RAW (Song et al., 2004) and THP-1 cells (Palacio et al., 2011). This observation may suggest a compromised immunoregulation of the inflammatory response of BV2 cells (Palacio et al., 2011). Another plausible explanation as to the reduced IL-10 expression in the presence of the hybrid protein and NAC is the fact that the microglia stimulated by Aβ have already differentiated to have an M-1 proinflammatory phenotype (Laffer et al., 2019) losing their ability to produce the anti-inflammatory IL-10.

The detection of secreted cytokine protein is by far the most widely used type of analysis. Because secreted protein is the biologically relevant form, its presence gives the most accurate picture of the cellular response (Sullivan et al., 2000). In addition, previous data show the absence of correlation between mRNA and protein expression levels of cytokines (Du et al., 2014; Israelsson et al., 2020; Maier et al., 2009). Hence, we no longer performed RT-PCR to quantify cytokine mRNA expression.

We have also demonstrated here that treatment of the hybrid protein significantly reduced the levels of both nitrite and cellular ROS in BV2 cells in a concentration dependent manner after Aβ treatment. Incidentally, a similar effect is also noted when BV2 cells were treated with LPS in the presence of the hybrid. A similar increase in inducible nitric oxide synthase (iNOS) expression leading to elevated levels of nitric oxide (NO) in glial cells during Aβ elicited inflammatory and immune responses has also been reported previously (Balez and Ooi, 2016). Aβ-RAGE interaction *in vitro* leads to cell stress with the generation of ROS and activation of downstream signaling mechanisms including the MAP kinase pathway (Yan et al., 2009).

Nonetheless, it can be noted that the levels of TNF-α, IL-1β, nitrite, and cellular ROS produced after oAβ stimulation are almost always greater compared to fAβ stimulation, consistent with the previous results that oAβ induces a more extensive inflammatory response and oxidative stress that cause greater damage in AD (Pan et al et al., 2011; Nirmalraj et al., 2020; Koffie et al.,2009; Lacor et al., 2007; Lambert et al., 1998; Pickett et al., 2016; Querol-Vilaseca et al., 2019; Shankar et al., 2008; Walsh et al., 2002; Wang et al., 2002). This reduction in the levels of the inflammatory cytokines TNF-α and IL-1β, nitrite and cellular ROS in the presence of the hybrid protein suggests that by diverting Aβ clearance from RAGE to the noninflammatory MerTK receptor, the activation of RAGE is also prevented. Because RAGE is not activated, the production of proinflammatory mediators (Behl, 2017; Ries and M. Sastre 2016) and microglia migration/infiltration which increases neuroinflammation (Wolfe, 2016; Dong et al., 2019) are reduced. The inhibition of RAGE that suppresses microglia activation and the associated inflammatory response and oxidative damage have also been previously documented (Yu and Ye, 2015; Fang et al., 2018; Hong et al., 2016).

Using the hybrid protein, MerTK-mediated phagocytosis of Aβ is promoted which may have led to downstream signaling that promotes ERK activation and the resolution of inflammation by promoting the synthesis of inflammation resolution mediators (Cai et al., 2018). This anti-inflammatory response may also involve a pathway that leads to the suppression of NF-kB-mediated signaling (Lee et al., 2012) or the inhibition of TLR-mediated innate immune response by activating STAT1, which selectively induces production of suppressors of cytokine signaling SOCS1 and SOCS3 (Lemke, 2013; Lemke and Rothlin, 2008; Rothlin et al., 2007; Zizzo and Cohen, 2018). Consistent with the previous results, the activation of MerTK by the hybrid protein also downregulates LPS-induced inflammation (Zhang et al., 2019).

Since the inflammatory and oxidative damage resulting from Aβ stimulation may in turn cause perturbations in the microglia’s ability for Aβ clearance or even stimulation of β-secretase for increased Aβ production (Hickman et al., 2008), the ability of the hybrid protein to not only facilitate Aβ clearance but also to reduce inflammation and oxidative stress through MerTK activation gives optimism for its potential as AD treatment and warrants the interrogation of its efficacy in *in vivo* testing. This result also highlights our novel strategy of developing the hybrid protein to target a specific ligand to a particular phagocytic receptor for the removal of Aβ. This approach can also be exploited for the therapeutic clearance of other deleterious metabolites by protein-guided phagocytosis-based approach.

Brain delivery is one of the major challenges in drug development because the BBB consists of capillary endothelial cells forming a restrictive barrier that selectively controls or prevents drugs from reaching their target (Zenaro et al., 2017; Jafari et al., 2019; Oller-Salvia et al., 2016; Zhou et al., 2021; Zlokovic, 1996). Here, we have shown that the hybrid protein crosses the BBB and colocalizes with both the microglia and MerTK receptor in the hippocampus 30 minutes after intraperitoneal injection. This result indicates that the APOE shuttle peptide that we used effectively mediated the delivery of the hybrid protein across the BBB by taking advantage of endogenous transport pathways (Oller-Salvia et al., 2016). This ability of the hybrid protein to cross the BBB is a necessary prerequisite to ensure putative target engagement in the brain when used *in vivo* for AD treatment.

The accumulation of Aβ peptide within the brain arises from an imbalance of the production and clearance of Aβ. Instead of the increased production of Aβ, recent evidence indicates that in most cases of AD, Aβ accumulation results because of an impaired clearance mechanism (Kurz and Perneczky, 2011; Mawuenyega et al., 2010; Ramanathan, et al., 2015; Wildsmith, et al., 2013). Hence, strategies that modulate Aβ clearance have been one important target in the development of disease modifying therapeutic agents for Alzheimer’s disease (Cummings et al., 2020; Cummings et al., 2018; Mawuenyega et al., 2010). The mechanisms of Aβ clearance and degradation include Aβ proteases, facilitated transport of Aβ across the BBB into the blood circulation (Deane et al., 2012), and uptake and phagocytosis of Aβ by cells such as microglia, astrocytes, and macrophages (Ries and Sastre, 2016).

After two months of treatment, we have demonstrated that the hybrid protein significantly reduced the levels of insoluble and total Aβ42 as well as the total Aβ42 to Aβ40 ratio in female double transgenic AD mice. In the brain, there are two main forms of Aβ. Under normal physiological conditions, the predominant Aβ species is 40 amino acids long (Aβ1-40) but in AD, there is an accumulation of Aβ1-42 (Bates et al., 2009) such that it becomes the principal species found in brain lysates of patients with AD (Nirmalraj et al., 2020). Moreover, Aβ42 has been shown to have higher tendency to aggregate and form fibrils (Nirmalraj et al., 2020; Murphy and LeVine, 2010; Bates et al, 2009; McGowan et al., 2005), and is considered as a key mediator of AD (Kwak et al., 2020).

It can be noted that the concentration of soluble Aβ42 in the brain is 20X lower compared to insoluble Aβ42. Interestingly, treatment of the hybrid protein for two months led to a significant reduction of almost 40% in insoluble Aβ42, the impact of which cannot be undermined because these insoluble dense fibrillar plaques act as reservoirs for the more toxic oligomers (Shankar et al., 2008; Haass and Selkoe, 2007). In studies of aducanumab, a monoclonal anti-amyloid antibody, removal of plaques was associated with a beneficial impact on cognitive decline, suggesting that fibrillar amyloid removal may correlate with disease modification in AD (Sevigny et al., 2016). Overall, the effect of the hybrid protein on the reduction of the ratio of total Aβ42/Aβ40 is more important because a high ratio is characteristic of many cases of known familial AD mutations showing that an increase in the Aβ42/Aβ40 ratio is important in AD pathogenesis. In addition, Aβ42/40 ratio and not the total Aβ level, plays a critical role in inducing neurofibrillary tangles in neurons (Kwak et al., 2020), a pathology that is more correlated with the severity of AD in humans.

Aβ in Alzheimer’s brain activates microglia and triggers a prolonged neuroinflammatory response that is tightly associated with synaptic dysfunction and cognitive decline (Zuroff et al., 2017; Wang et al., 2015). In turn, neuroinflammation aggravates Aβ deposition suggesting positive feedback between aggregation and inflammation (Hughes et al., 2020). These activated microglia play a potentially detrimental role by eliciting the expression of proinflammatory cytokines such as IL-1β, IL-6, and TNF-α influencing the surrounding brain tissue (Wang et al., 2015; Saito and Saido, 2018). Experimental and clinical evidence demonstrate the increased synthesis of proinflammatory cytokines such as TNF-α, IFN-γ, IL-1β, IL-6, IL-18, and the upregulation of their cognate receptors in the AD brain (Wang et al., 2015).

Here, we have shown that hybrid protein treatment reduces the levels of IL-6 in the brain of female double transgenic AD mice. IL-6 is a pleiotropic inflammatory cytokine mainly produced by activated microglia and astrocytes capable of stimulating these cells to release a cascade of other proinflammatory cytokines. The levels of IL-6 have been found significantly elevated in the brains, cerebrospinal fluid, and plasma, especially locally around amyloid plaques in AD patients and animal models (Wang et al., 2015; Cojocaru et al., 2011; Hüll et al., 1996; Wu et al., 2015). IL-6 induces calcium influx in the hippocampal neurons through N-methyl-D-aspartate (NMDA) receptors (Orellana et al., 2005) which has been demonstrated to trigger synaptic collapse (Arbel-Ornath et al., 2017).

IL-6 in neuroinflammation and neurodegeneration plays a complex role in regulating cognitive function where elevated IL-6 is correlated with Aβ aggregation and the appearance of hyperphosphorylated tau in AD brain (Chen et al., 2012) as well as with age-related cognitive decline in humans (Wang et al., 2015). IL-6 is also considered a useful biological marker to correlate with the severity of cognitive impairment in AD (Lai et al., 2017). Considering the strong association between peripheral IL-6 and cognitive decline (Bradburn et al., 2018), its reduction in the brain after two months of treatment suggests the potential of the hybrid protein in mitigating the production of this proinflammatory cytokine, thereby regulating the neuroinflammatory environment and cognitive decline in AD.

The observed reduction in Aβ burden in the brain and IL-6 concentration in the brain also resulted to better survival of the nerve cells. According to Cummings (2017), neuroprotection might be achieved by direct effects on neurons (primary neuroprotection) or by interfering with processes that lead to cell death such as inflammation. Cell survival may be achieved through interruption of upstream processes leading to cell death or through affecting downstream processes that lead to neuronal death. In AD, removal of amyloid plaques has been demonstrated using several immunotherapies to have no corresponding clinical benefit and reducing fibrillar amyloid demonstrated with amyloid imaging is not disease modifying (Cummings, 2017; Holmes et al., 2008; Liu et al., 2015). However, in animal models, the reduction of amyloid plaque by AVCRI175 therapy was accompanied by a reduction of neuroinflammation and oxidative damage in the brain of APP/PS1 mice (Muñoz-Torrero, 2017). Most likely, the neuroprotection resulting from hybrid protein treatment is due to the reduction of IL-6, a potent and damaging inflammatory cytokine in AD, which might have been the offshoot of the reduction in Aβ reduction.

We have also shown that the liver and kidney histology did not manifest histopathological abnormalities characteristic of injury due to drug toxicity. This result implies that the hybrid protein is safe as it did not cause any toxicity even after two months of daily treatment at 50 mg/kg body weight. Because the liver is responsible for concentrating and metabolizing any drug, it is a prime target for drug-induced damage which may be the result of direct toxicity from the administered drug or their metabolites, or injury may result from immune-mediated mechanisms (David and Hamilton, 2010). Some examples of liver histopathological abnormalities associated with drug treatment include hematopoiesis, necrosis, karyomegaly, cytomegaly, intranuclear acidophilic cytoplasmic invagination, inflammation, and fibrosis (Dorso et al., 2016).

On the other hand, the kidneys play an essential role in preserving homeostasis of the body’s internal environment, including regulation of water, electrolyte, nitrogen, and acid–base balances. Hence, toxic drugs may impair renal function and cause drug-induced nephrotoxicity (Faria et al., 2019). Examples of kidney histopathological abnormalities associated with drug treatment include basophilic tubules, karyomegaly and cytomegaly, proteinaceous casts, and glomerulosclerosis (Dorso et al., 2016). While different blood biomarkers are available to assess both liver (Church and Watkins, 2017; Kullak-Ublick et al., 2017) and kidney (Faria et al., 2019; Gobe et al., 2015; Griffin et al., 2019) damage associated with drug treatment, we performed H and E staining to visualize possible hepatic, glomerular, and medullary damage that may result from treatment of the hybrid protein.

Taken together, our results in both cellular and animal models of AD provide a proof of concept showing the utility of the hybrid protein to specifically channel the clearance of Aβ to MerTK. Though the primary AD target of the hybrid protein is Aβ clearance, its specificity to MerTK receptor endows it with the secondary AD mitigating capability by reducing inflammation, thus conferring a certain level of neuroprotection in the double transgenic mouse model of AD. Thus, these findings show the potential of the hybrid protein as AD treatment that can target both the Aβ deposition and inflammation aspects of the disease.

The costs of caring for AD patients was estimated to have reached $1 trillion globally in 2018 and if a cure will not be developed, financial burden is projected to rise to $2 trillion by 2030 (Cummings et al., 2020). Hence, the development of a disease-modifying treatment or a cure is urgently needed.

The disease pathology of AD is still poorly understood. However, the three main hallmarks are Aβ plaque deposition, tau tangle formation, and massive neuroinflammation resulting to the death of nerve cells (Dong et al., 2019; Zenaro et al., 2017; Doens and Fernandez, 2014). Recent efforts towards AD treatment focus on the mechanisms that may decrease or alter the dynamics of accumulation of the above proteins, and some neuroprotective interventions (Cummings et al. 2019). While a wide array of molecular targets and potential therapeutic compounds have been identified in the past 15 years, most of these targets were found ineffective in randomized clinical trials (Cummings et al. 2020). There are only currently a few anti-Aβ agents that have shown positive clinical or biomarker effects in long-duration trials and pharmacological characteristics. The most recently approved agent for AD was aducanumab, a monoclonal antibody that partially targets oligomers, while mostly clearing insoluble amyloid plaques (Cummings et al. 2020). Other agents with the potential for near term approval are the injectable antibodies, gantenerumab, BAN2401, and a small molecule oral agent, ALZ-801 (see Cummings et al 2020 for review). Gantenerumab acts similarly with aducanumab; while BAN2401 preferentially targets soluble protofibrils (large oligomers) over plaques; and ALZ-801 blocks the formation of oligomers without binding to plaques. Several immunotherapeutic approaches, such as anti-Aβ and anti-pyroglutamate antibodies are also ongoing (Hettman et al 2020; Sperling et al 2011). However, these antibodies form immune complexes that can trigger the complement cascade, activate perivascular macrophages, and induce vasogenic edema and cerebral microhemorrhages with these anti-Aβ therapies (Hettman et al 2020; Sperling et al 2011). Moreover, the risk of adverse events in response to anti-Aβ monoclonal antibody therapy appears to be greater in apolipoprotein E4 carriers (Sperling et al 2011), which is the major population being targeted.

The Aβ clearance facilitated by hybrid protein through the noninflammatory pathway offers an attractive alternative for potential treatment. In the absence of formation of immune complexes, it may significantly reduce the adverse events associated with most currently available anti-Aβ immunotherapeutic approaches, particularly the risk of vasogenic edema and cerebral microhemorrhages, with antibody-mediated clearance of Aβ.

## MATERIALS AND METHODS

### Cell culture

BV2 mouse microglial cells (RRID: CVCL_0182) were grown in Dulbecco’s Modified Eagle Medium (DMEM) with 4.5 g/L glucose (Cytiva #SH30243.02) supplemented with 10% fetal bovine serum (GE Life Sciences #SH30910.03), 2 mM L-Glutamine (GE Life Sciences #SH30852.01), 100 units/mL Penicillin and 100 μg/mL Streptomycin (Corning #30-001-CI) in a standard tissue culture incubator with 5% carbon dioxide, unless otherwise specified.

### Preparation of Aβ42

We prepared Aβ1-42 oligomers (oAβ) and fibrils (fAβ) following Evans (2006) and Ryan et al (2010) using siliconized polypropylene tubes (VWR #4165SL). We used Aβ1-42 to test the effects of the hybrid protein because it is more predominant in AD plaques (Nirmalraj et al., 2020) and its level is higher compared to the other Aβ species in the blood plasma of AD patients (Butterfield et al., 2013; Izzo et al., 2014; Verdier and Penke, 2004). Briefly, synthetic Aβ1-42 (Anaspec AS-60479-01) was prepared by resuspending the lyophilized protein in hexafluoroisopropanol (HFIP MPBio #151245). After overnight air drying to completely remove HFIP, the Aβ film was stored at -20°C. Immediately prior to use, Aβ was resuspended in dimethyl sulfoxide (DMSO #WN182) and sonicated for 10 minutes. The oligomers were prepared by diluting the DMSO stock with serum-free and phenol red-free DMEM (GE Life Sciences #SH30284.01) and incubating at 4°C for 24 hours without shaking. The fibrils were similarly prepared but incubated at 37°C for 24 hours with vigorous shaking.

### Protein expression and purification

The hybrid protein and AβBP were cloned into pMAL-C5E MBP vector (New England Biolabs) with a cleavage site for 3C protease immediately after MBP. The fusion proteins were expressed in BL21 (DE3) *E. coli,* purified with amylose (New England Biolabs #E8022L) columns, eluted with maltose (VWR #1B1184), and dialyzed against 1X PBS. On the other hand, the MPD- containing Tubby N-terminal was expressed and purified as described previously (Caberoy et al., 2010a). The purified FLAG-tagged proteins were cleaved from MBP or Glutathione S-transferase (GST) using 3C protease (Genscript #Z03092-100) and their purity was analyzed by SDS-PAGE.

### Phagocytosis assay

BV2 cells were seeded at 30-40% confluency (1.5-2 X 10^5^ cells per mL) onto 24-well plates (Corning #3524) with round cover slips (VWR #89015-725) at the bottom a day before the assay. One hour before treatment, the cell culture media was changed into serum-free and phenol red-free DMEM. Phagocytosis assays were done by incubating BV2 cells with 100 *µ*M HiLyte™ Fluor 488-labeled human Aβ1-42 (Anaspec #AS-60479-01) for three hours in the presence or absence of the hybrid protein at 37°C for Aβ uptake and at 4°C for surface binding assay following Prasad and Rao (2018).

### Immunofluorescence and confocal microscopy

Phagocytosis assays were performed for three hours. BV2 cells were then washed with ice cold DPBS (Cytiva #SH30028.02) three times and fixed in 4% paraformaldehyde (Thermo Fisher #J61899-AP) for 3 minutes. Cells were then washed three times in DPBS for three minutes, quenched in 50mM NH_4_Cl for five minutes, washed again, and permeabilized in 0.1% Triton X-100 for 30 minutes at room temperature. Blocking was done using Intercept blocking buffer (Licor #927-70001) for one hour. Primary antibodies (α-FLAG Sigma-Aldrich Cat# F1804, RRID:AB_262044, α-MerTK Santa Cruz Biotechnology Cat# sc-365499 AF647, RRID:AB_10843860, α-non-muscle myosin IIA BioLegend Cat# 909801, RRID:AB_2565100) were incubated overnight at 4°C, followed by a repeated wash in PBS-T, first every five minutes for an hour, and again every 10 minutes for an additional hour. Secondary antibodies (α-mouse Alexa Fluor 568 Molecular Probes Cat# A-11004, RRID:AB_2534072) at 1:300 dilution were incubated for one hour at room temperature, repeatedly washed in PBS-T first every five minutes for two hours, and again every 10 minutes for an additional two hours. The cells were then mounted using Vectashield mounting medium with DAPI (Vector Labs #H-1200). Confocal images were taken using Nikon Air1 at 40X magnification for quantification, and at 100X magnification for the representative photomicrographs. The images were taken as maximum projections of z-stacks at 1*μ*m per stack. Analysis was performed using ImageJ Fiji software (Fiji, RRID:SCR_002285), a Java-based image processing program developed at the National Institutes of Health and the Laboratory for Optical and Computational Instrumentation. Fluorescence data of Aβ associated with BV2 cells were normalized to the fluorescence of cells incubated without the hybrid protein, following background subtraction using cells not fed with Aβ.

The brain of a 15-month-old double transgenic AD mouse was fixed in 4% paraformaldehyde for 4 hours at 4°C or for 1 hour at room temperature. The fixed brain was then washed with 1X PBS and dehydrated in increasing sucrose concentration (5%, 15%, and 30%) until it sank to the bottom of the container. Infiltration was done overnight at 4°C using 50% Optimal Cutting Temperature (OCT Tissue Tek #4583) and 50% of 30% sucrose with gentle shaking followed by imbedding in 100% OCT. Thioflavin T (ThT Chem-Impex #22870) staining was done to verify the presence of Aβ plaques in the brain of APP/PS1 AD mouse. Coronal sections at 10 *µ*m thickness was permeabilized for 1 hour using 1% Triton X in 1X PBS. Permeabilization was followed by staining with 0.08% ThT in 50% ethanol for seven minutes. Differentiation was done for 10 seconds each in decreasing concentrations of ethanol from 100%, 95% and 90% followed by washing in 1X PBS for four hours. The stained sections were then mounted in Fluoromount (Electron Microscopy Sciences #17984-25) with DAPI for confocal microscopy.

### Fluorescence activated cell sorting analysis

Phagocytosis assay and time course clearance were carried out as described above. BV2 cells were then harvested and fixed in 0.5% paraformaldehyde overnight at 4°C. The cells were then centrifuged at 1500 X g for 5 minutes and subsequently washed with 1X DPBS. Cells were filtered through a 40 *µ*m cell strainer (VWR #10199-655) to ensure single cell suspension and analyzed using Sony SH800 flow cytometer at 488/535 nm excitation/emission. The samples were run at a low flow rate and either 30,000 or 100,000 events were collected. Similar to confocal analysis, fluorescence data of Aβ associated with BV2 cells were normalized to the fluorescence of cells incubated without the hybrid protein, following background subtraction using cells not fed with Aβ. Quantification of fluorescence intensity and the generation of histogram and density plots were done using the Sony SH800 analysis software.

### Cell survival or cytotoxicity assay

The effects of the hybrid protein on the survival of BV2 cells during Aβ clearance was assessed using Cell Counting Kit - 8 (CCK-8 Dojindo #CK04), following the manufacturer’s protocol. This colorimetric assay is based on the principle of a change in color of water-soluble tetrazolium (WST-8 - (2-(2-methoxy-4-nitrophenyl)-3-(4-nitrophenyl)-5-(2,4-disulfophenyl)-2H- tetrazolium, monosodium salt) to orange as formazan is formed upon reduction of WST-8 by cellular dehydrogenase. WST tetrazolium salt was used because its detection sensitivity is higher than any other tetrazolium salts such as MTT (3-(4,5-dimethylthiazol-2-yl)-2,5-diphenyltetrazolium bromide), XTT (2,3-Bis(2-methoxy-4-nitro-5-sulfophenyl)-2H-tetrazolium-5-carboxanilide), or MTS (3-(4,5-dimethylthiazol-2-yl)-5-(3-carboxymethoxyphenyl)-2-(4-sulfophenyl)-2H- tetrazolium).

Briefly, 5,000 cells after three hours phagocytosis assay in the presence of 200 nM hybrid protein were dispensed in a 96-well plate (Corning #3596) at a total volume of 100 *µ*L in serum-free and phenol red-free DMEM. After 3, 6, 12, 24, 48, and 72 hours incubation at 37°C, 5% CO2, 10 *µ*L of CCK-8 solution was added to each well. Absorbance at 450 nm was measured following three hours incubation using a Spectra Max Plus plate reader.

### Competition and blocking studies

To test for the specificity of hybrid protein-mediated phagocytosis to MerTK, we used protein molecules to compete with or block the binding of the hybrid protein to the cell surface receptor. To compete with MerTK and RAGE receptors, we added an excessive amount (1:50 dilution) of the extracellular domains of MerTK (Mer-Fc R and D Systems #891-MR) and RAGE (RAGE-Fc R and D Systems #1145-RG), respectively to the hybrid protein and Aβ 30 minutes prior to their incubation with BV2 cells. The idea is that Mer-Fc, but not RAGE-Fc will bind to the MPD of the hybrid protein therefore preventing it from directing the clearance of Aβ through MerTK. Dominant negative blocking was done by adding 200 nM of AβBP6 to Aβ 30 minutes prior to incubation with the hybrid protein and BV2 cells. Similarly, Aβ will bind to AβBP6, but since the AβBP6 has no MDP that can recognize MerTK, it will not direct Aβ clearance through MerTK. To block any interactions with the ligand binding domains of the receptors, antibodies that bind specifically MerTK (Abcam #ab184086 or FabGennix #MKT-101AP) and RAGE (R and D Systems Cat# AF1179, RRID:AB_354649) were added at 1:200 dilution 30 minutes prior to feeding the cells with Aβ in the presence of the hybrid protein.

### *Ex vivo* phagocytosis

To test whether the hybrid protein can also facilitate the uptake of physiologically generated Aβ, we did *ex vivo* phagocytosis assay following Borhmann et al. (2012) using sections of AD mouse brain previously verified to have Aβ plaques. To differentiate the live BV2 cells from the endogenous mouse microglial cells present in the brain sections, we used GFP-expressing cells as effectors. BV2 cells were transfected with GFP plasmid using TransIT transfection reagent (Mirus #MIR5404) following the manufacturer’s protocol. Forty-eight hours after transfection, BV2 cells were harvested by trypsinization and seeded onto 10 cm plate with double transgenic AD mouse brain sections mounted in a glass slide. After three hours incubation at 37°C, the slide was processed for immunohistochemistry (IHC) and confocal microscopy, as described above. Aβ was detected using clone 6E10 Aβ-specific antibody (BioLegend Cat# 803017, RRID:AB_2565327) and α-mouse Alexa Fluor 568 (Molecular Probes Cat# A-11004, RRID:AB_2534072) secondary antibody.

### Multiplex ELISA and confocal microscopy for cytokines

BV2 cells were cultured in 6-well plate (Greiner Bio #657 160). Upon reaching 80% confluency, the cells were treated with 1 *µ*m non-fluorescent Aβ1-42 (Anaspec #AS-20276) in the presence or absence of the hybrid protein in phenol red-free and serum-free DMEM. After 24 hours incubation at 37°C, 5% CO_2_, the media were collected, and cellular debris was removed by centrifugation at 3,000 X g for 5 minutes. The supernatant was stored at -80°C until analysis. The concentration of secreted cytokines was quantified by multiplex ELISA using the mouse high sensitivity T-Cell discover array 18-plex (MDHSTC18) of EveTechnologies. LPS (Enzo #ALX-581-019-L002) at 1 *µ*g/mL and NAC (Millipore Sigma #194603) at 1 mM were used as reference. 200 nM TubN and AβBP6, peptides that contain part of the hybrid protein, were included as controls.

Complementary to ELISA, intracellular cytokines were also detected by immunofluorescence microscopy following (Freer and Rindi, 2013) with some modifications. Briefly, BV2 cells were incubated in 1 *µ*M non-fluorescent Aβ for two hours then an EZCell™ Protein Transport Inhibitor Cocktail (BioVision #K932-500) which disrupts the structure and function of the Golgi apparatus and allows cytokines to be retained intracellularly, was added. Cells were incubated for an additional four hours followed by immunofluorescence processing for confocal microscopy described previously. Intracellular cytokines were detected using the following primary antibodies: anti-TNF-α (Thermo Fisher Scientific Cat# AMC3012, RRID:AB_2536391), anti-IL-1β (Abcam Cat# ab9722, RRID:AB_308765), anti-IL-6 (Abcam Cat# ab6672, RRID:AB_2127460) and anti-IL-10 (Bioss #bs-069812) followed by anti-rabbit IgG Alexa Fluor 488 (Thermo Fisher Scientific Cat# A-11034, RRID:AB_257621). Images were transformed using ImageJ Fiji software (Fiji, RRID:SCR_00228).

### Greiss assay to determine nitrite concentration

To test the effect of the hybrid protein on the production of nitric oxide by BV2 cells upon Aβ stimulation, we looked at the level of its stable metabolic product, nitrite using Greiss assay. Briefly, BV2 cells were cultured in a 24-well plate (Corning #3524) with 1 *µ*m non-fluorescent Aβ in the presence or absence of the hybrid protein using serum-free and phenol red-free DMEM. After 24 hours incubation at 37°C, 5% CO_2_, the media were collected, and cellular debris was removed by centrifugation at 500 X g for 5 minutes. Colorimetric assay was done using Greiss reagent (Enzo #89150-572) following the manufacturer’s protocol. Absorbance at 540 nm was taken using a Spectra Max Plux plate reader. LPS at 1 *µ*g/mL and NAC at 1 mM were used as positive and negative controls, respectively.

### DCFDA (dichlorofluorescin diacetate) assay for cellular reactive oxygen species ROS

Cellular reactive oxygen species (ROS) was quantified using 2’,7’– dichlorofluorescin diacetate (DCFDA) assay. This assay is based on diffusion of DCFDA into the cell which is then deacetylated by cellular esterases to a non-fluorescent compound, which is later oxidized by ROS into a highly fluorescent 2’,7’–dichlorofluorescein (DCF). Briefly, BV2 cells were seeded onto black flat-bottomed 96-well plate (Sycamore #655086). The culture media were changed to serum-free and phenol red-free DMEM one hour before treatment with 1 *µ*M non-fluorescent Aβ in the presence or absence of the hybrid protein. After 24 hours incubation at 37°C and 5% CO_2_, the media were removed and replaced with 150 *µ*L of 20 *µ*M DCFDA (Millipore Sigma #287810) in DPBS. Cells were incubated at 37°C for one hour and protected from light until quantification of fluorescence using FLX800 microplate fluorescent reader at Ex/Em 485/535 nm. LPS at 1 *µ*g/mL and 1 mM NAC, a known antioxidant, were used as controls.

### Animals

All animal protocols were approved by the Institutional Animal Care and Use Committee (IACUC) of the University of Nevada, Las Vegas. The APP/PS1 double transgenic AD mice (B6.Cg-Tg(APPswe,PSEN1dE9)85Dbo/Mmjax; Stock Number: 034832-JAX; RRID:MMRRC_034832-JAX) purchased from the Jackson Laboratory (Bar Harbor, Maryland) were housed and bred under standard conditions in groups of three to six per cage. To assess whether the hybrid protein can cross the BBB, 200 mg/kg body weight of the hybrid protein was administered intraperitoneally, and the mice were sacrificed after 30 minutes. To determine if the hybrid can ameliorate AD, six-month-old female mice (25.2-29.8g BW) were randomly assigned to treatments. Six replicates for each treatment were used. Hybrid protein (50 mg/kg BW) was administered by intraperitoneal injection once daily on alternating sides on progressive days of administration. Insulin needles were used to avoid injury (perforations) to the bowel. The control group was injected with PBS. After two months, mice were sacrificed and blood and tissue collections were done. The left brain was fixed for IHC analysis while the right brain, together with the other organs, was frozen at -80°C for biochemical analysis.

### Tissue processing and immunohistochemistry (IHC)

The left brain was fixed in 4% parafolmaldehyde for 4 hours at 4°C or for 1 hour at room temperature. The fixed organ was then washed with 1X PBS and dehydrated in increasing sucrose concentration (5%, 15%, and 30%) until it sank to the bottom of the container. Infiltration was done overnight at 4°C using 50% Optimal Cutting Temperature (OCT Tissue Tek #4583) and 50% of 30% sucrose with gentle shaking followed by imbedding in 100% OCT. Coronal sections of 10 *µ*m thickness along the hippocampus were used for IHC (described previously). Detection of Aβ, neurons, microglia, astrocytes, MerTK, and the hybrid protein in the brain was done using these antibodies: α-Aβ (Biolegend clone 6E10 #803017), α-MAP2 (ThermoFisher #MA5-12826), α-IBA1 (Abcam #ab178846), α-GFAP (Thermo Fisher #13-0300), α -MerTK (Sta Cruz #sc-365499 AF647), and α-FLAG (Sigma #F1804) and the appropriate secondary antibodies (Invitrogen anti-rabbit IgG Alexa Fluor 488 #A-11034, 647 #A-21244; anti-mouse IgG Alexa Fluor 488 #A-10680, 568 #A-11004).

The same tissue processing was done using the liver and kidney for hematoxylin (Ricca #3536-1) and eosin (Electron Microscopy #26763-03) staining except that the kidney was cut at 20-22 *µ*m thickness. Images were obtained using the Axio Zeiss inverted compound light microscope and Zen lite 2012 Zeiss imaging software.

### Biochemical analyses

Cytokine concentration in the brain of mice was determined using a Multi-Analyte ELISArray kit (Qiagen) following the manufacturer’s protocol. Both the positive and negative controls were provided in the kit. The levels of Aβ40 and Aβ42 were quantified by sandwich ELISA using Aβ40 and Aβ42 EZbrain ELISA kit (EMD Millipore) following the manufacturer’s protocol.

### Statistical analysis

Statistical significance at P ≤ 0.05 between treatments, each with three replicates unless otherwise specified, was analyzed using two-tailed Student’s t-test. Data are expressed as mean ± SEM.

### Data availability statement

The original contributions presented in the study are included in the article/supplementary materials. Further inquiries can be directed to the corresponding author.

## Acknowledgments

The authors thank the other members of the laboratory for technical assistance. This work was partially supported by NIH grants K99EY020865 / R00EY020865 (NBC) and P20GM121325 (NBC) and a UNLV TTDGRA to LPS. Confocal imaging was performed at the UNLV Confocal and Biological Imaging Core with the assistance of Sophie Choe. The authors would also like to thank Casey Hall of UNLV Genomics Core Facility, Linda Loverro of the Animal facility, Shirley Shen of the Nevada Institute of Personalized Medicine Core Laboratory, and Mimi Zhang and Dr. Anthony Shen of the Urban Air Quality Laboratory for their assistance with DNA sequencing, flow cytometry, and fluorescent plate reader, respectively.

## Authors contributions

NBC, LPS, PP and AS contributed to the study conception and design. Material preparation and data collection were performed by LPS, PP, AS, KE, DU, PE, KC, JAMG,AE, NR, and NBC. Data analyses and writing of the manuscript were done by LPS and NBC. All authors read and commented on the previous version of the manuscript and approved the submitted version.

## Supplementary Figures

**Supplementary Figure 1.**
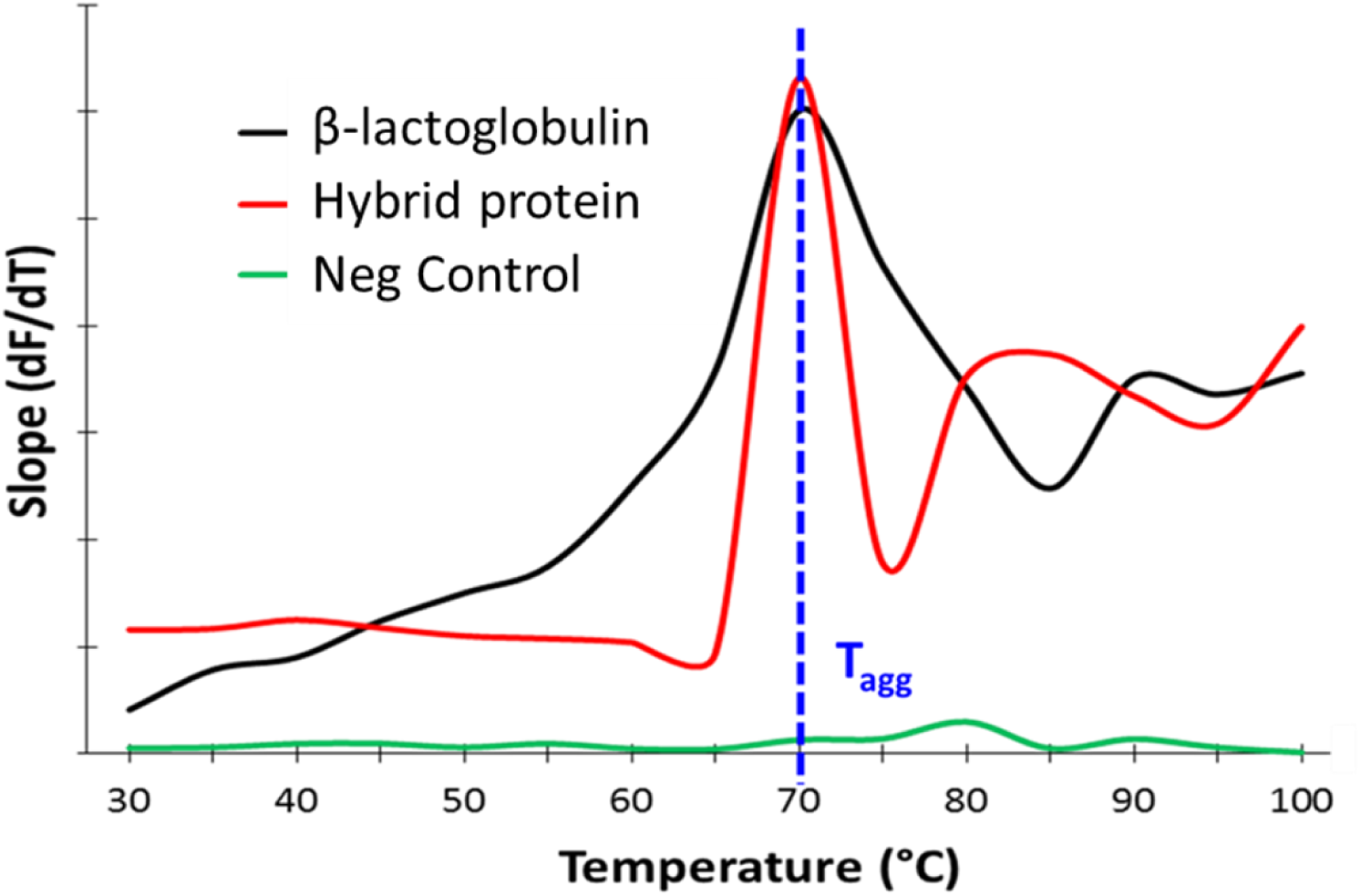
The Hybrid is stable at physiological conditions. The stability of nAβBPh was determined using PROTEOSTAT® Thermal Shift Stability Assay Kit (Enzo Life Sciences). We assayed 8 mg/mL hybrid protein at pH7 following the manufacturer’s protocol. β-lactoglobulin and buffer only were used as positive and negative controls, respectively. Note that at the concentration and pH we used, nAβBPh was relatively stable at physiological conditions, and started to aggregate only at 70°C.

**Supplementary Figure 2.**
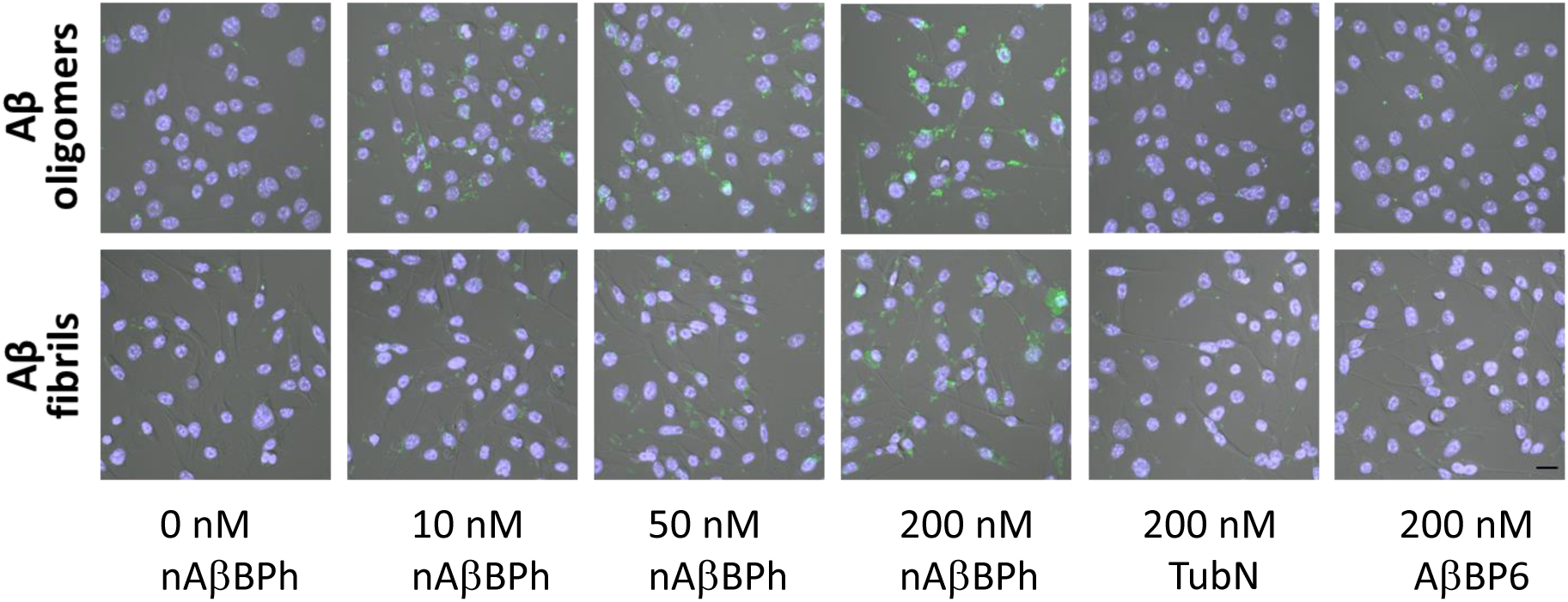
related to Figure 2. Confocal analysis shows that the hybrid protein facilitates significantly higher phagocytic efficiency of Aβ by BV2 cells. Representative confocal images of BV2 cells for the quantification of internalized Aβ per cell and percentage of phagocytic cells. The fluorescence intensity and exposure settings were kept constant. Fluorescence data were normalized to the negative control (0 nM nAβBPh) following background subtraction using cells not fed with Aβ. Scale bar = 20 μm. n = 10 fields with at least 20 cells/field.

**Supplementary Figure 3.**
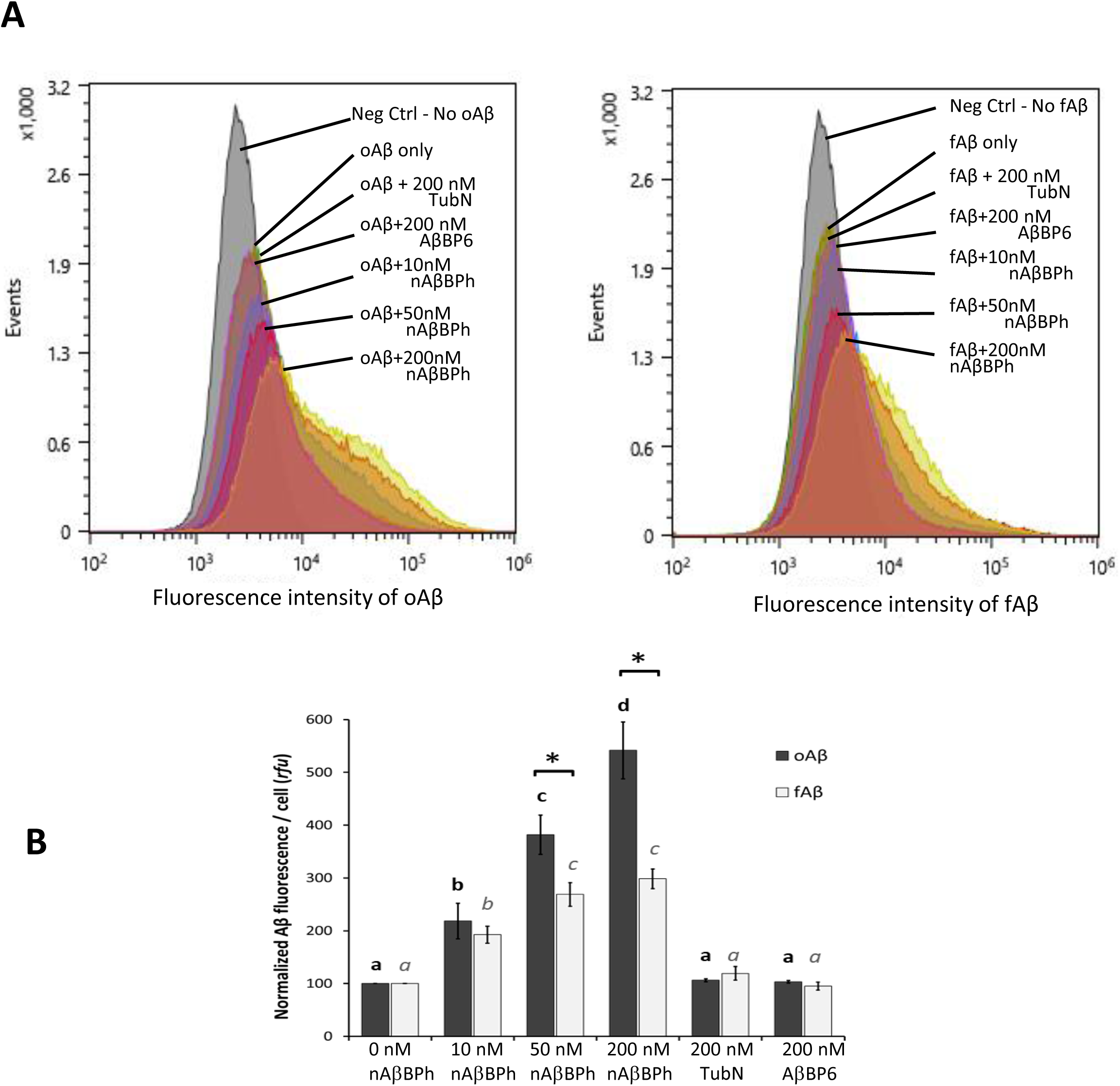
FACS analysis shows that the hybrid protein facilitates significantly higher uptake of Aβ by BV2 cells. (A) Representative histograms for oAβ (left) and fAβ (right). (B) Quantification demonstrates the effects of the presence of the hybrid protein on cell-associated Aβ after three hours phagocytosis assay. The X-axis shows Aβ fluorescence in logarithmic scale, and the Y-axis represents the number of cells. n = 3 independent experiments with at least 30,000 cells per treatment. Different letters denote statistical significance at P ≤ 0.05 between treatments using unpaired, two-tailed Student’s t-test. Asterisk (*) denotes significant differences between the means of oAβ and fAβ in the same treatment. Data are expressed as mean ± SEM.

**Supplementary Figure 4.**
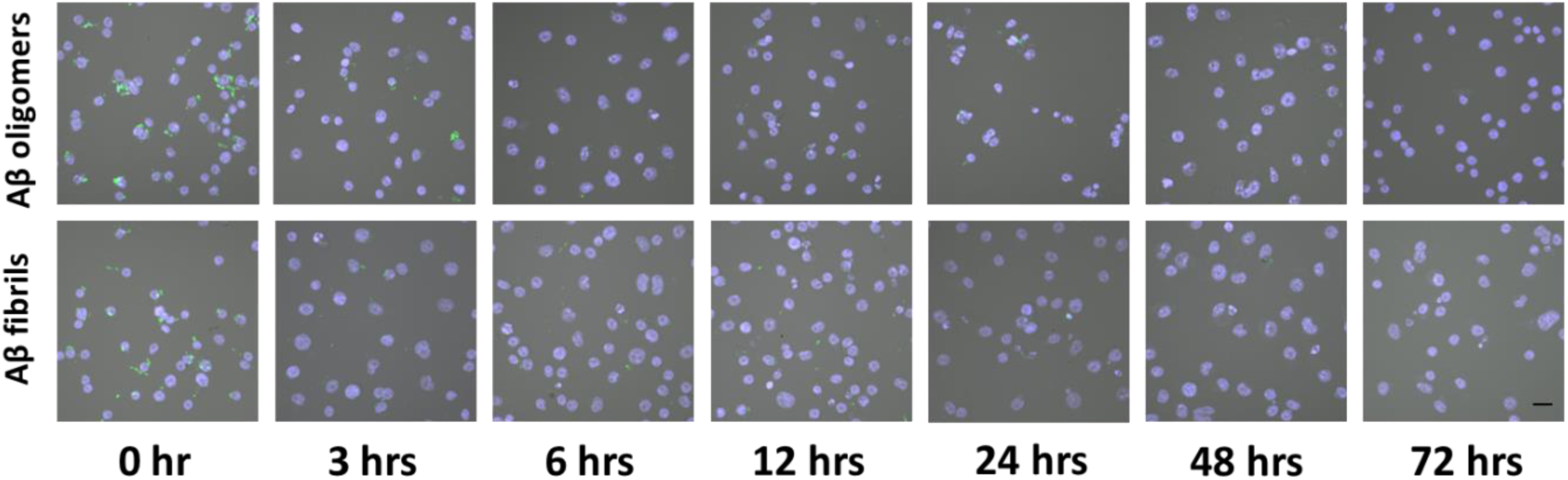
related to **Figure 4.** BV2 cells effectively clears Aβ phagocytosed in the presence of the hybrid protein. Representative confocal images of BV2 cells for the quantification of cell associated Aβ in a time course. The fluorescence intensity and exposure settings were kept constant. Fluorescence data were normalized to oAβ at 0 hour following background subtraction using cells not fed with Aβ. Scale bar = 20 μm. n = 18 fields with at least 20 cells/field.

**Supplementary Figure 5.**
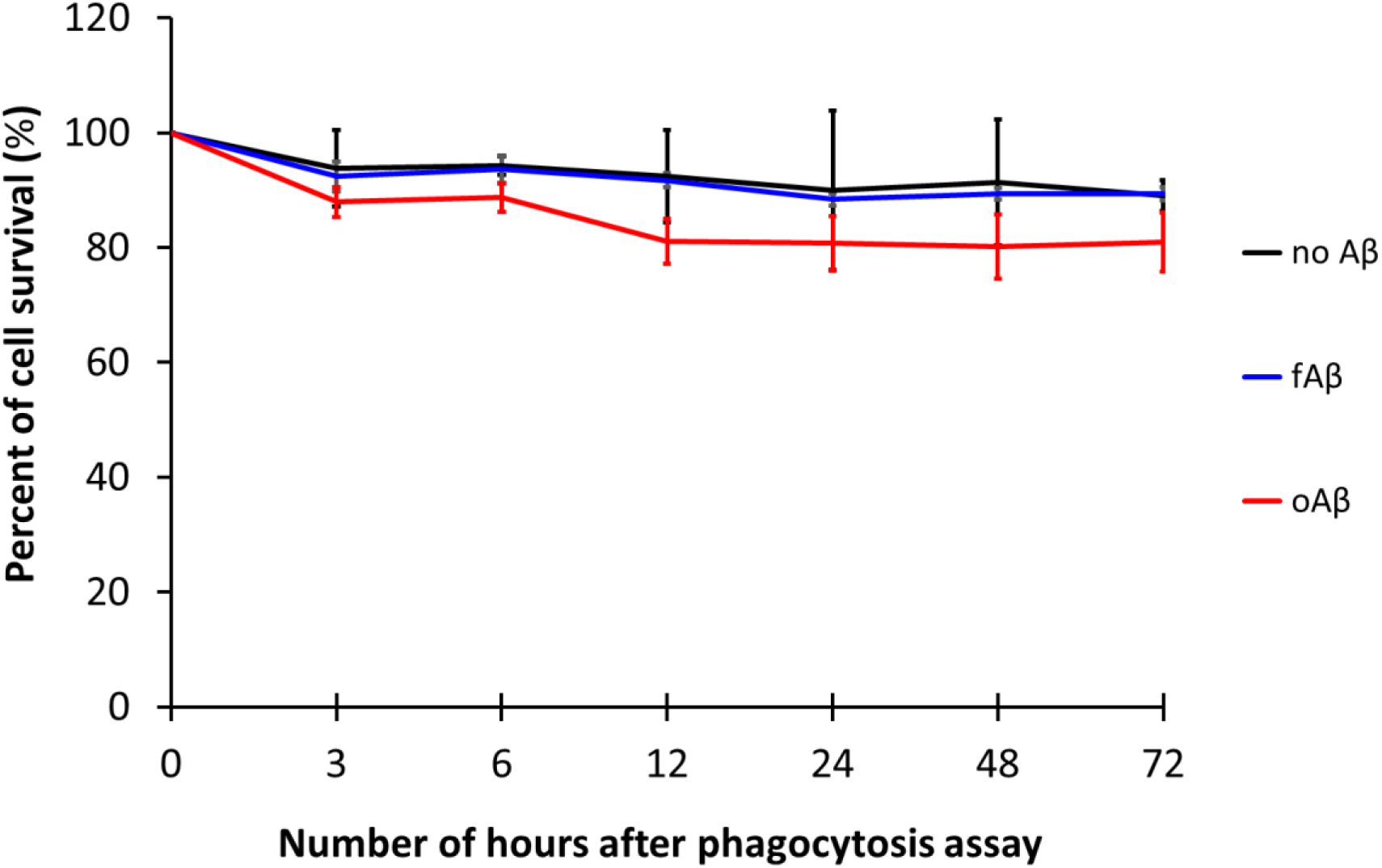
Phagocytic clearance of Aβ has no significant effect on the survival of BV2 microglial cells. Viability of the cells in the presence or absence of the hybrid protein using Water Soluble Tetrazolium (WST)-8-based colorimetric assay. Cell survival and cytotoxicity assay was determined using Cell Counting Kit – 8 following the manufacturer’s protocol. n = 4 independent experiments with 3 replicates each. P ≤ 0.05. Data are expressed as mean ± SEM.

**Supplementary Figure 6.**
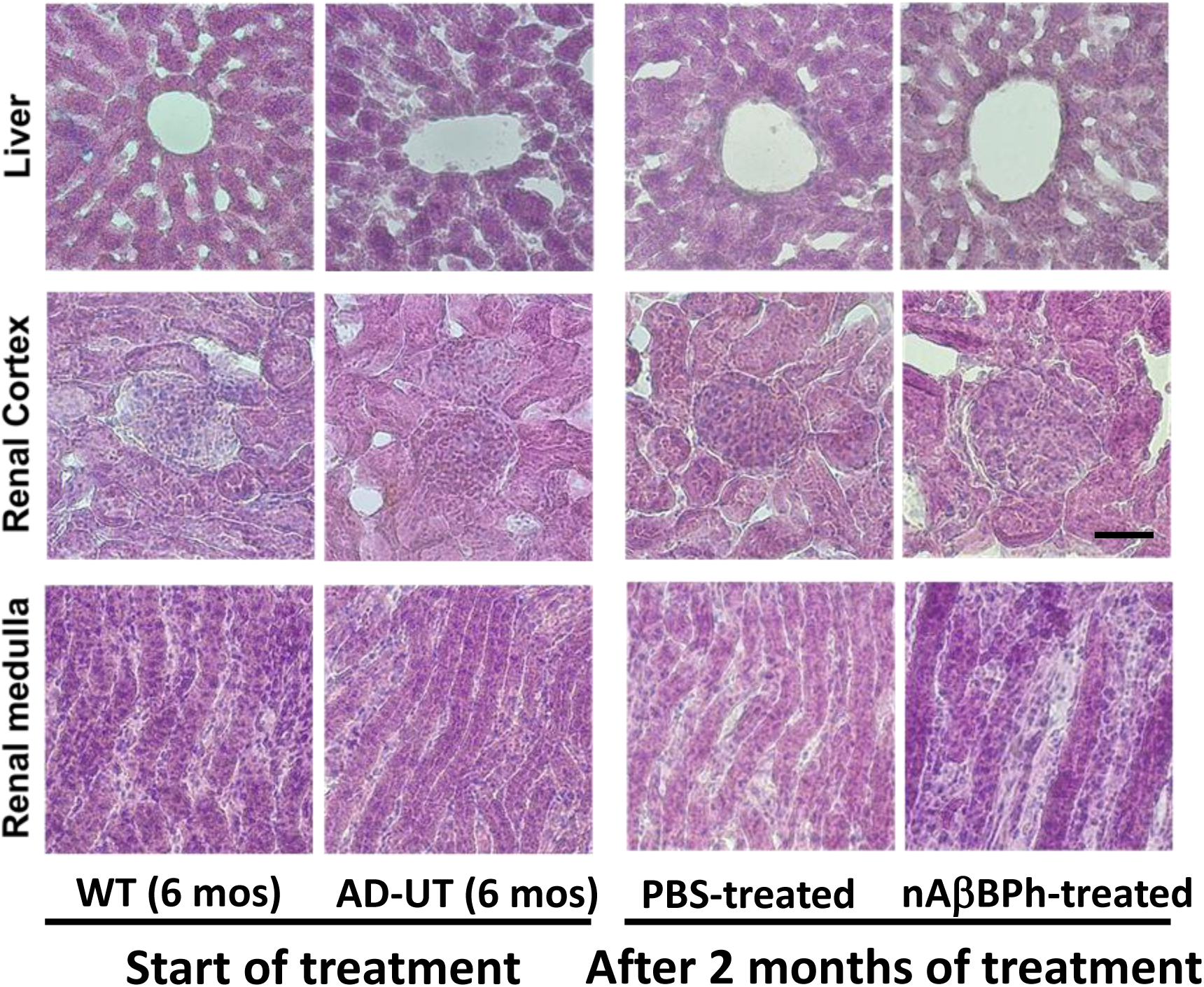
Hybrid treatment does not cause histopathological toxicity to the liver and kidney. Representative H and E staining of the liver, and the renal cortex and renal medulla of the kidney show normal histology even after one month of daily injection of the hybrid protein. AD-UT= untreated APP/PS1. n = 6 mice. Scale bar = 50 *µ*m.

